## Supplementary material 2: Pre workshop questions for "Addressing cultural and knowledge barriers to enable preclinical sex inclusive research": Supplementary material 2 - Pre workshop questions.pdf

### Exploring researchers' perspective on sex inclusive in vivo research Pre-workshop survey

If you have any questions about this survey, please find a survey facilitator or contact Prof. Amrita Ahluwalia or Dr. Jonathan Ho

* Required

1. By leaving your initials below, you acknowledge that you have had the opportunity to read the survey information sheet and consent form and have the research study explained. You have had the opportunity to ask questions about the research study, and your questions have been answered. You understand that if you are uncomfortable with any question, you may skip it.

   **Please make sure to write the same initials (minimum 3) below when filling out the pre and post surveys.** *

2. Does your current research use animal models to investigate your disease of interest? *

- ◯ Yes
- ◯ No

3. Does your research focus on a disease, or phenomena, that can occur in both sexes? *

- ◯ Yes
- ◯ Sometimes
- ◯ No

4. How often are you involved or can influence the planning of experiments involving animals? *

- ◯ Never
- ◯ Rarely
- ◯ Sometimes
- ◯ Often
- ◯ Always

5. In the past, how much statistical training have you received?

- ◯ No training
- ◯ Primarily informal/practical training (e.g., books, videos, discussion)
- ◯ 1-2 formal courses (e.g., Coursera, University class)
- ◯ More than 2 courses or a statistical degree

6. How familiar are you with factorial experimental designs (please consider your general knowledge and application of the technique)?

- ◯ Not at all familiar
- ◯ Not very familiar
- ◯ Somewhat familiar
- ◯ Familiar
- ◯ Extremely familiar

7. Thinking over the last 5 years, how often have you incorporated both sexes into your experiment while studying an intervention (for example a drug treatment)?

- ◯ Never: 0% of the time
- ◯ Rarely: 1-25% of the time
- ◯ Sometimes: 26-50% of the time
- ◯ Often: 51-75% of the time
- ◯ Almost always: 76-100% of the time

**Over the next few questions, answer based on your current thinking about how incorporating sex into an in vivo experimental design may affect the overall design, data or statistical analysis resulting from that study.**

8. Do you think inclusion of both sexes requires doubling a study's sample size?

   - ◯ Yes
   - ◯ Sometimes
   - ◯ No

9. Do you think sex influences data variability, therefore when you include both sexes, more animals are needed?

   - ◯ Yes
   - ◯ Sometimes
   - ◯ No

10. When analyzing *in vivo* data collected from both sexes, do you think sex should be included in the statistical model?

- ◯ Yes
- ◯ Sometimes
- ◯ No

11. When analyzing *in vivo* data collected from both sexes, do you think the analysis should be disaggregated (for example run a t-test comparing control and intervention on the male data and then running a separate test on the data from the females)?

- ◯ Yes
- ◯ Sometimes
- ◯ No

12. When analyzing *in vivo* data, do you think data from the two sexes should be pooled (combined) for the analysis (for example run a single t-test comparing control and intervention group)?

- ◯ Yes
- ◯ Sometimes
- ◯ No

**Below we ask questions pertaining to <u>including both sexes in an *in vivo* experiment in addition to the intervention of interest.</u> This means that both sexes are planned into the experiment at the beginning stages. However, the purpose of the study does not have to include specific questions pertaining to the differences between the sexes.**

13. Overall, I think that **including both sexes into an *in vivo* experimental design** is....
    1= Bad 7= Good

| 1 | 2 | 3 | 4 | 5 | 6 | 7 |
|---|---|---|---|---|---|---|

14. Overall, I think that **including both sexes into an *in vivo* experimental design** is....
    1= Worthless 7= Useful

| 1 | 2 | 3 | 4 | 5 | 6 | 7 |
|---|---|---|---|---|---|---|

15. Overall, I think that **including both sexes into an *in vivo* experimental design** is....
    1= Harmful 7= Beneficial

| 1 | 2 | 3 | 4 | 5 | 6 | 7 |
|---|---|---|---|---|---|---|

16. Overall, I think that **including both sexes into an *in vivo* experimental design** is....
    1= The wrong thing to do 7= The right thing to do

| 1 | 2 | 3 | 4 | 5 | 6 | 7 |
|---|---|---|---|---|---|---|

17. I feel confident in my ability to **include both sexes into an *in vivo* experimental design**

- ◯ Strongly disagree
- ◯ Disagree
- ◯ Somewhat disagree
- ◯ Neither agree nor disagree
- ◯ Somewhat agree
- ◯ Agree
- ◯ Strongly agree

18. Whether or not I **include both sexes in an *in vivo* experimental design** is completely up to me

- ◯ Strongly disagree
- ◯ Disagree
- ◯ Somewhat disagree
- ◯ Neither agree nor disagree
- ◯ Somewhat agree
- ◯ Agree
- ◯ Strongly agree

19. Overall, using **both sexes in an *in vivo* experimental design** is

- ◯ Extremely difficult
- ◯ Moderately difficult
- ◯ Slightly difficult
- ◯ Neither easy nor difficult
- ◯ Slightly easy
- ◯ Moderately easy
- ◯ Extremely easy

20. In my professional environment, stakeholders I respect think I should **include both sexes into an *in vivo* experimental design**

- ◯ Strongly disagree
- ◯ Disagree
- ◯ Somewhat disagree
- ◯ Neither agree nor disagree
- ◯ Somewhat agree
- ◯ Agree
- ◯ Strongly agree

21. I feel professional pressure from my scientific community to **include both sexes into an *in vivo* experimental design**

- Strongly disagree
- Disagree
- Somewhat disagree
- Neither agree nor disagree
- Somewhat agree
- Agree
- Strongly agree

22. It is expected of me that I **include both sexes into an *in vivo* experimental design**

- Strongly disagree
- Disagree
- Somewhat disagree
- Neither agree nor disagree
- Somewhat agree
- Agree
- Strongly agree

23. I expect to **include both sexes into an *in vivo* experimental design** in my next in vivo experiment

- Strongly disagree
- Disagree
- Somewhat disagree
- Neither agree nor disagree
- Somewhat agree
- Agree
- Strongly agree

24. I want to **include both sexes into an *in vivo* experimental design** in my next *in vivo* experiment

- Strongly disagree
- Disagree
- Somewhat disagree
- Neither agree nor disagree
- Somewhat agree
- Agree
- Strongly agree

25. I intend to **include both sexes into an *in vivo* experimental design** in my next *in vivo* experiment

- ◯ Strongly disagree
- ◯ Disagree
- ◯ Somewhat disagree
- ◯ Neither agree nor disagree
- ◯ Somewhat agree
- ◯ Agree
- ◯ Strongly agree

26. Select which of the following prevents you from **including both sexes** in future experiments in addition to the intervention of interest (choose **all** that apply):

- ☐ Availability of sample/test material
- ☐ Complexity of experimental design
- ☐ Convention
- ☐ Cost
- ☐ Data analysis concerns
- ☐ Experiment would take longer
- ☐ Female animals are more variable
- ☐ Logistics
- ☐ Male animals are more likely to fight and may lead to premature euthanasia
- ☐ Model behavior may be different in the other sex
- ☐ Not relevant to the research question
- ☐ Sample size concerns
- ☐ Welfare issues
- ☐ Other

27. Select which of the following you believe are the advantages, if any, to **including both sexes in an in vivo experimental design**? (choose **all** that apply):

- ☐ 3Rs – Reduction
- ☐ Animal welfare
- ☐ Efficient use of all animals from breeding
- ☐ Reproducibility
- ☐ Translatability
- ☐ Understanding sex differences
- ☐ Other

28. Do you have any other thoughts or comments you would like to provide regarding **including both sexes in an in vivo experimental design**?

29. What is your age?

Number must be between 18 ~ 110

30. Which of the following best describes your gender? (If you would prefer to self-describe, please do so in the free-text box.)

- Man
- Non-binary
- Woman
- Prefer not to say
- Other

31. How many years have you worked with animals in research?

Number must be between 0 ~ 70

32. What is the primary type of *in vivo* research in which you are involved (if several, chose the option you identify most with)?

- ○ Basic biological investigation
- ○ Discovery (e.g., target identification)
- ○ Drug development/Lead optimization
- ○ Production quality and manufacturing
- ○ Research support (e.g., statisticians, ethical review board, or animal technician)
- ○ Safety pharmacology/preclinical safety/toxicology
- ○ Other

33. What is your highest level of education?

- ○ Doctoral degree (PhD, veterinary degree, other)
- ○ Master's degree
- ○ Bachelor's degree (4 years at college/university)
- ○ Associate or vocational qualification (2 years at college/university)
- ○ High school diploma, GED, A level, or equivalent
- ○ Other

34. If you’d like to be entered into the draw for a £50 Amazon gift card, please leave your email address below

This content is neither created nor endorsed by Microsoft. The data you submit will be sent to the form owner.

Microsoft Forms
