## Supplementary material 4 - workshop training material for "Addressing cultural and knowledge barriers to enable preclinical sex inclusive research": Supplementary material 4 - workshop training material.pdf

### Workshop: Best Practice for Sex Inclusive Research

1: Quantitative Biology, Discovery Science, R&D, AstraZeneca, UK.

2:  Queen Mary University of London, UK.

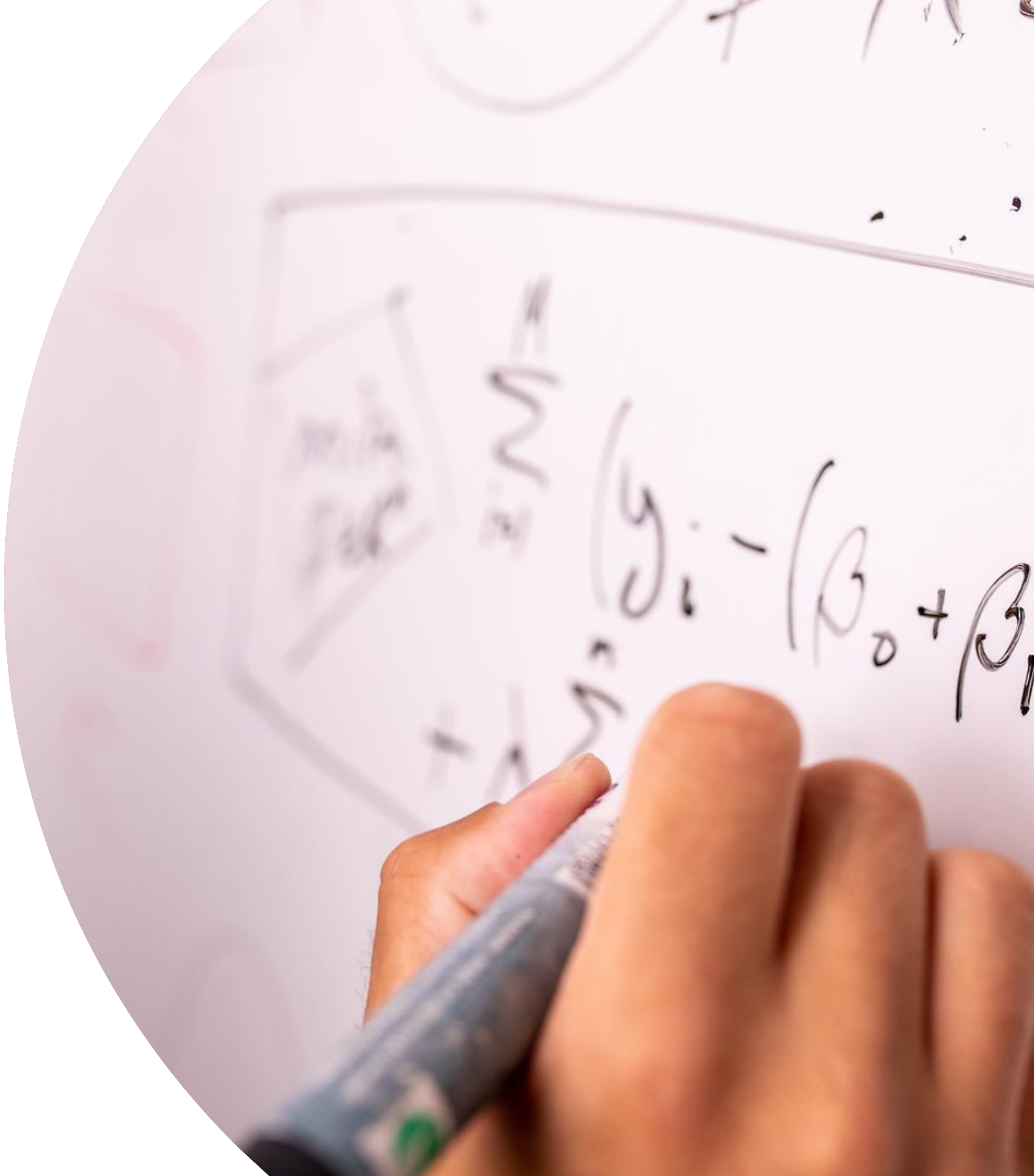

### Outline

1 Terminology

2 Clinically sex matters

3 What is happening in preclinical research?

4 What is sex inclusive research?

5 Factorial analysis is critical element

6 Considering the barriers

7 Conclusion

1

### Terminology

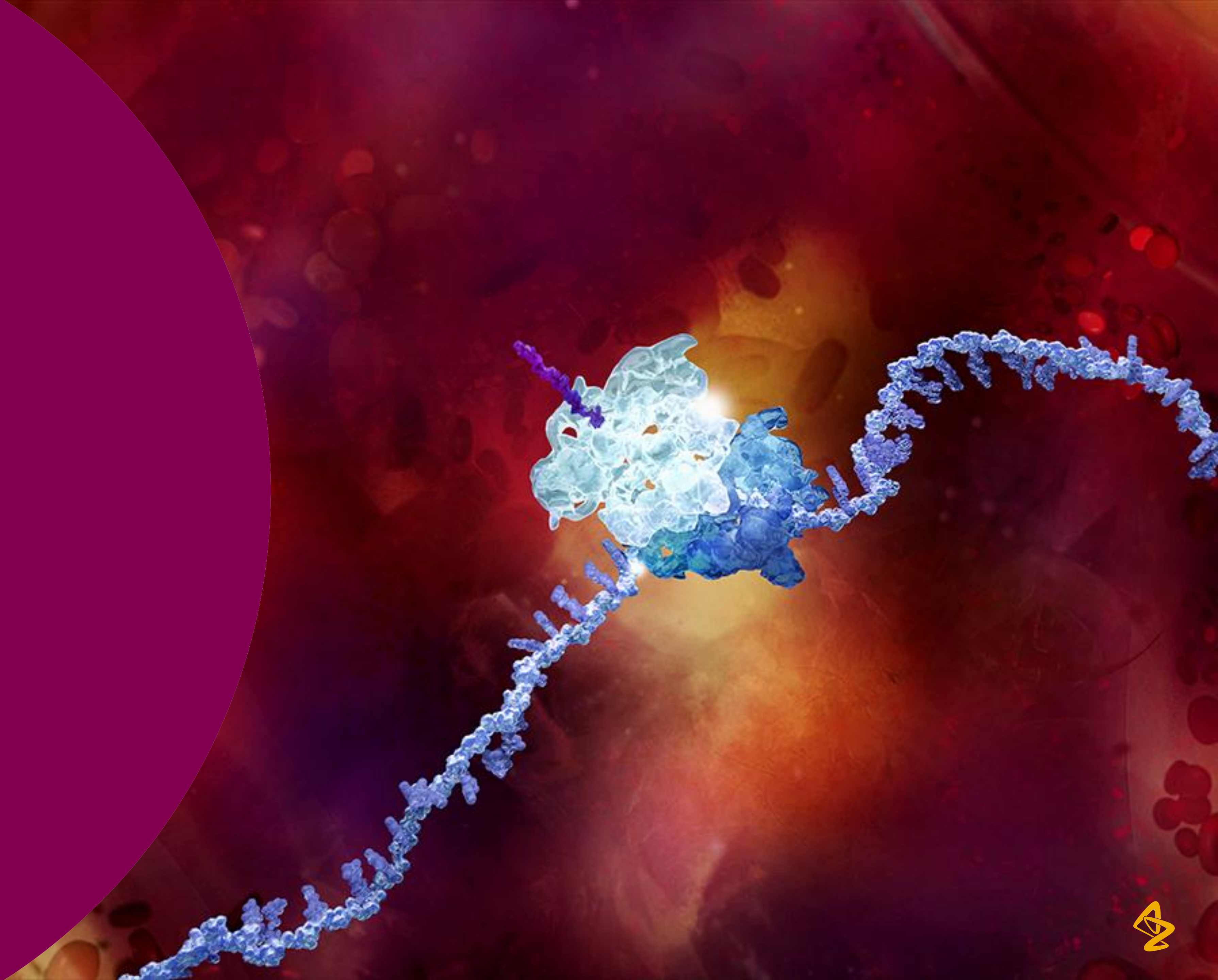

### Gender / Sex – does terminology matter?

**Sex refers to**

- *"the different biological and physiological characteristics of males and females, such as reproductive organs, chromosomes, hormones, etc"* *World Health Organisation*

**Gender refers to**

- *"the socially constructed characteristics of women and men – such as norms, roles and relationships of and between groups of women and men. It varies from society to society and can be changed."* *World Health Organisation*

2

### Clinically does sex matters?

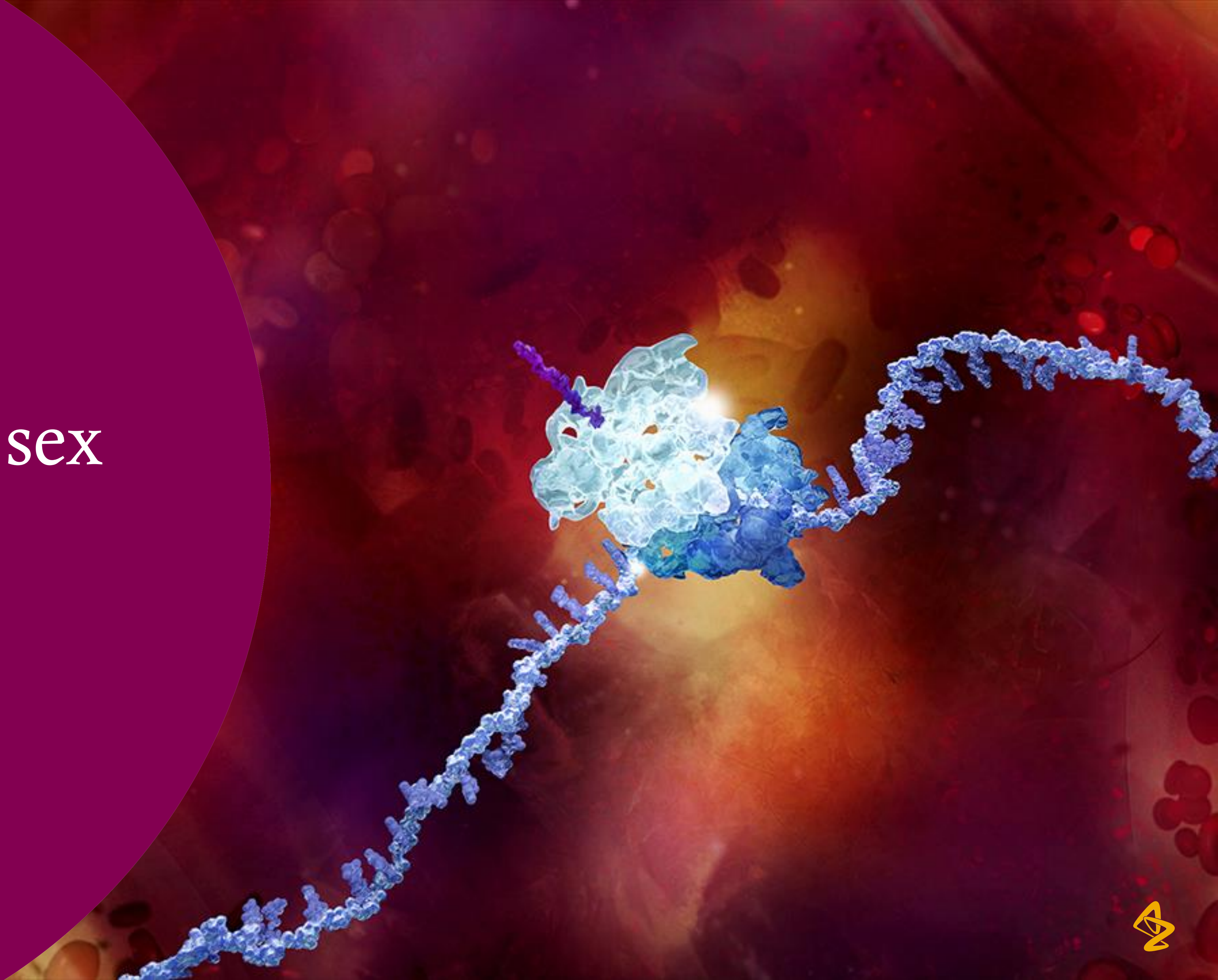

### Sex matters clinically

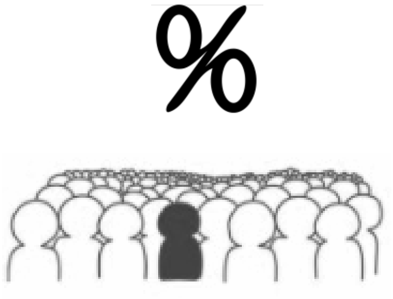


Prevalence

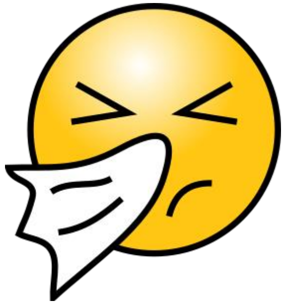

Symptoms

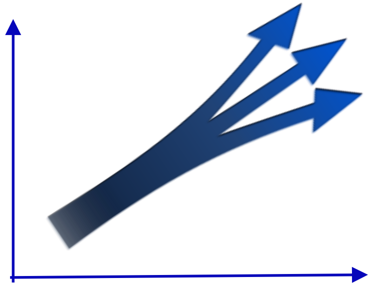

Progression

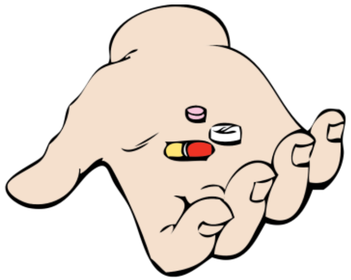

Side Effects

**COVID-19** [Bwire 2020; Doerre & Doblhammer 2022]

- Prevalence higher in ♀ but higher morbidity and mortality in ♂
  - Biological differences ?
    - Higher expression ACE 2 receptor for coronavirus in ♂
    - Immunological differences driven by sex hormone and X chromosome
- Gender differences
  - ♀ - more contacts, work in care roles
  - ♂ higher rates of smoking and drinking
  - ♂ Lower uptake of preventative measures

3

### What is happening in preclinical research?

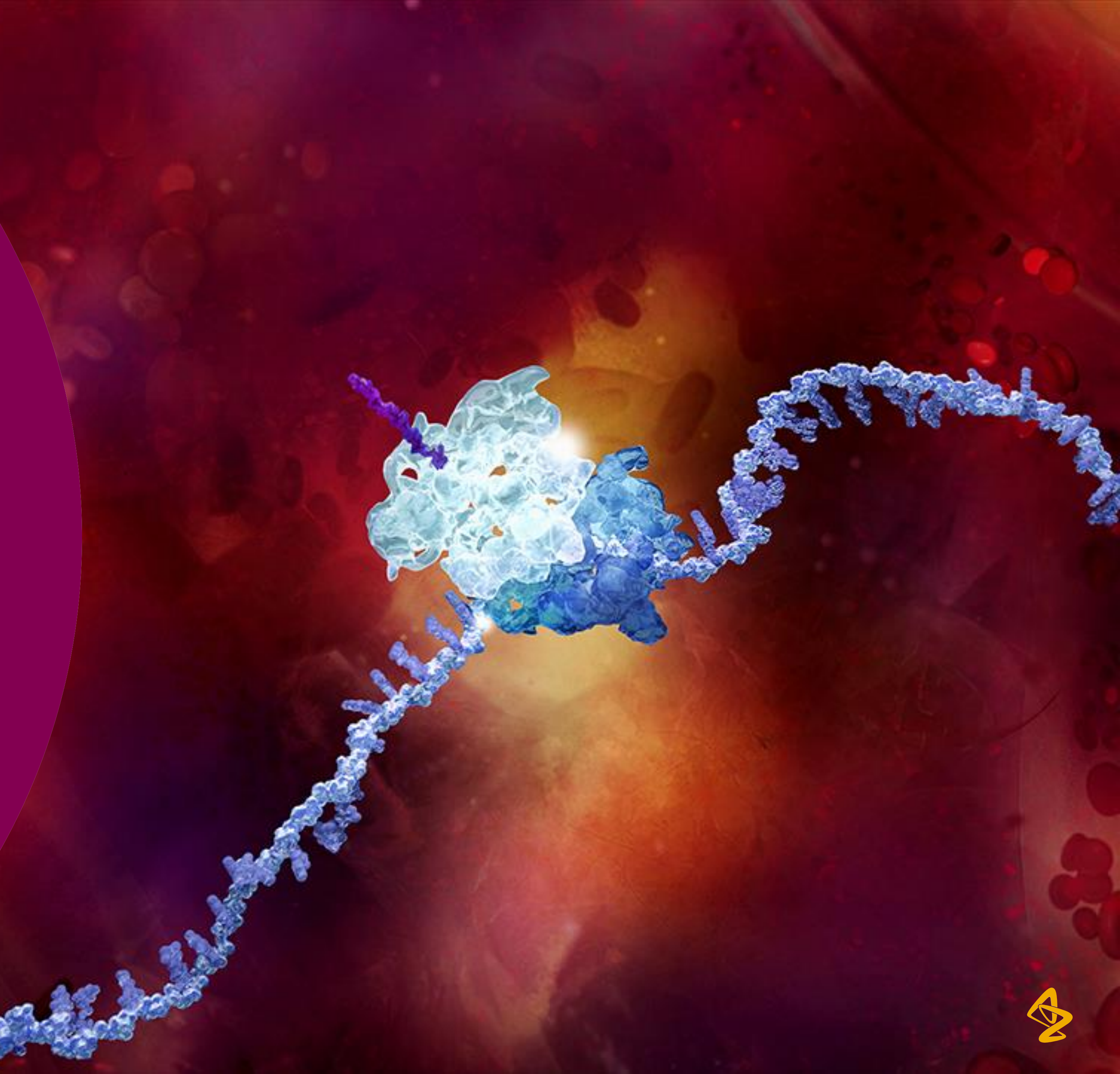

### Classic design - driven by minimising N

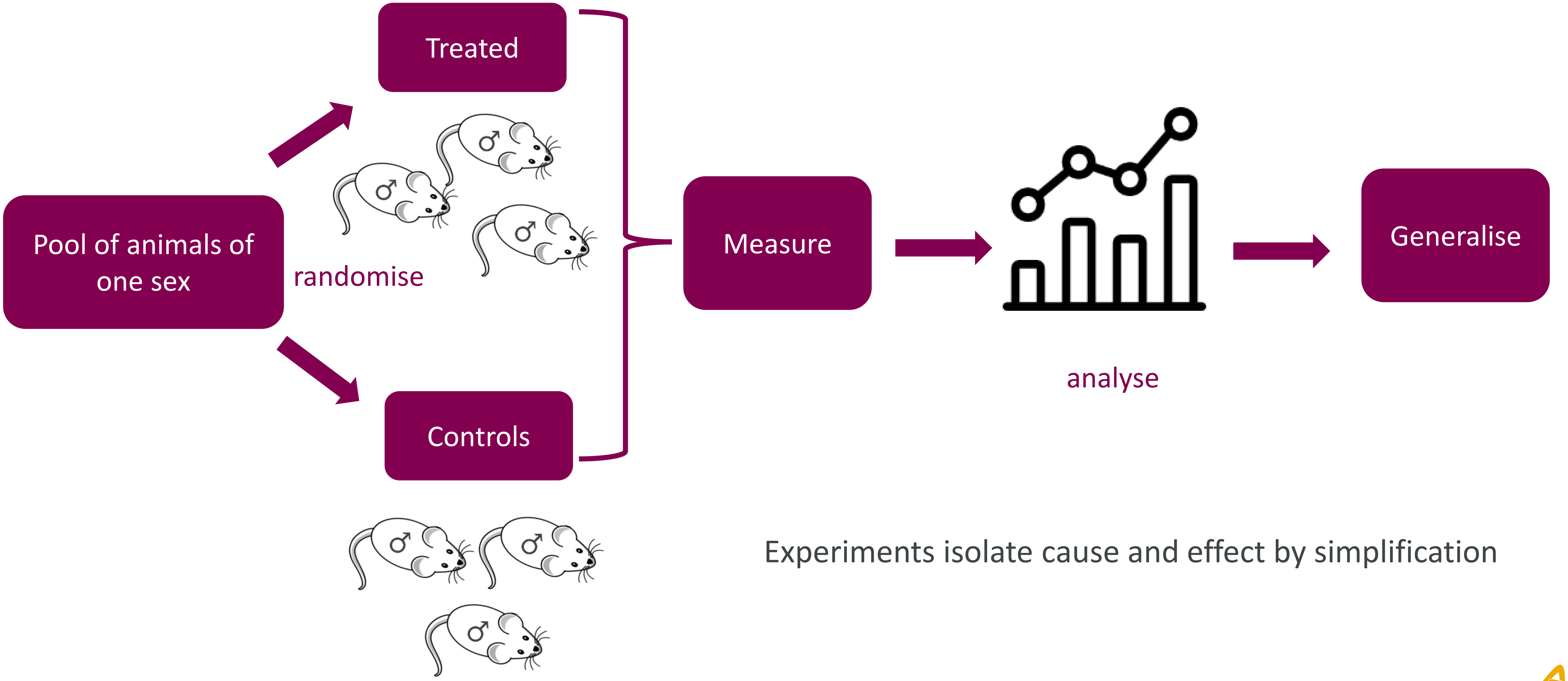


Experiments isolate cause and effect by simplification

### Embedded neglect of sex within preclinical research

- Reporting:
  - In vivo: Sex not specified – 22% did not specify Yoon et al 2014
  - In vitro: 75% did not report the sex Shah 2014
- Experimental design:
  - In vivo:  comparison across 9 fields of biology, 2009 to 2019 Beery 2020
    - 6/9  significant improvement,  1(Pharmacology) reduction to 29%, average 26% to 48%
  - In vitro: 69 -80% male only Taylor 2011, Shah 2014
- Analysis (In vivo):
  - When both sexes (N=356), only 42%  sex-based analysis Beery 2020
  - Those reporting sex differences:  1/3 did not test  statistically
  - Garcia-Sifuentes & Maney 2021

### Emerging evidence that our knowledge base is biased

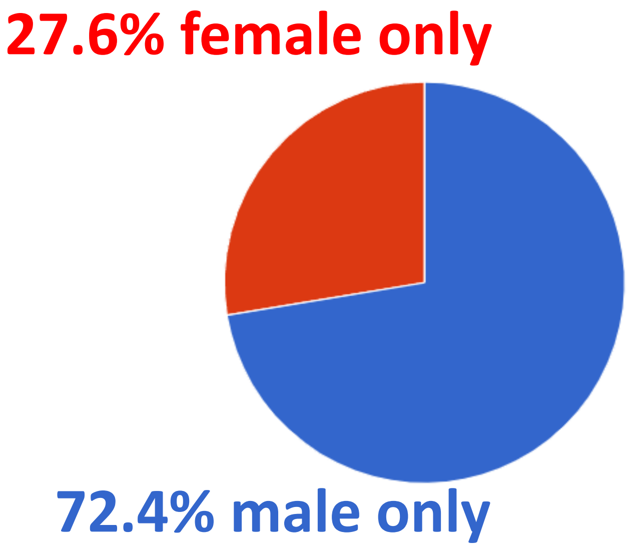


Pain processing

N=127

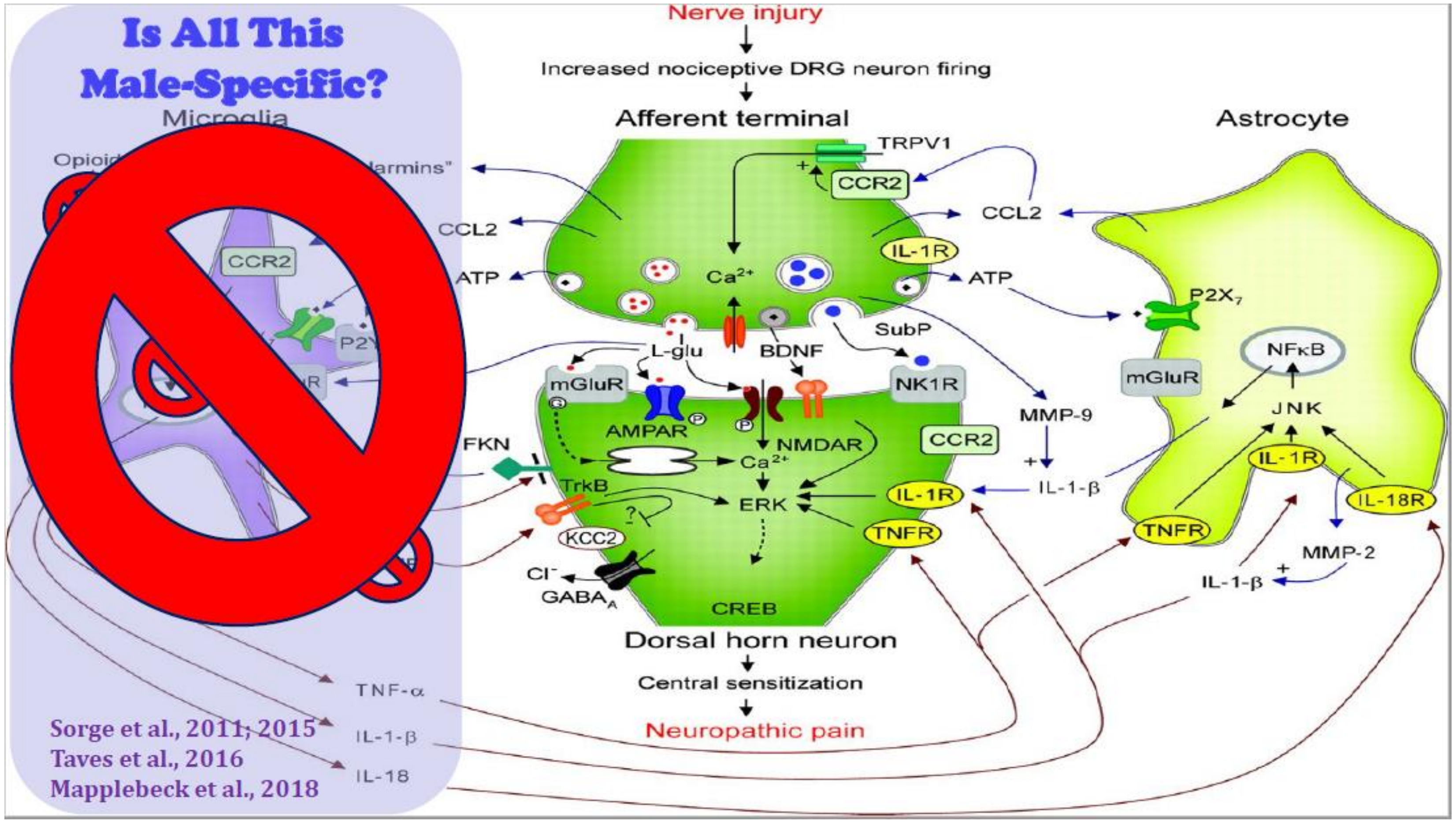


Mogil (2020) Nature Reviews Neuroscience

Figure courtesy of Prof Jeffery Mogil (McGill University Canada) 2022

### Evidence: sex matters in early research

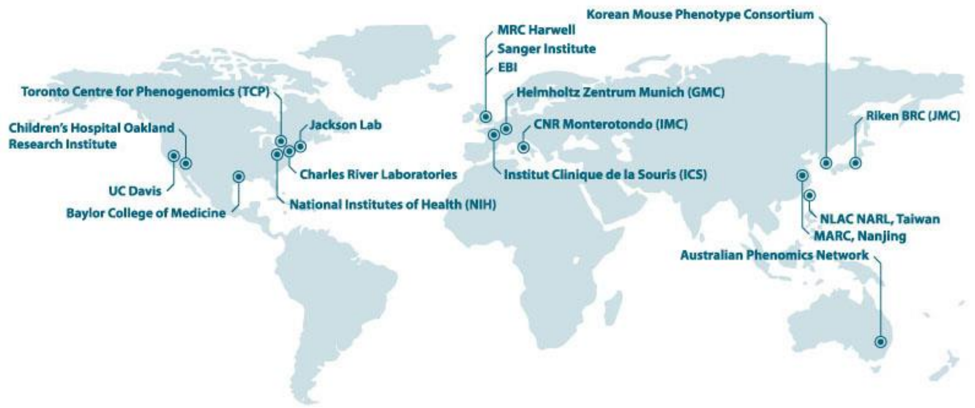


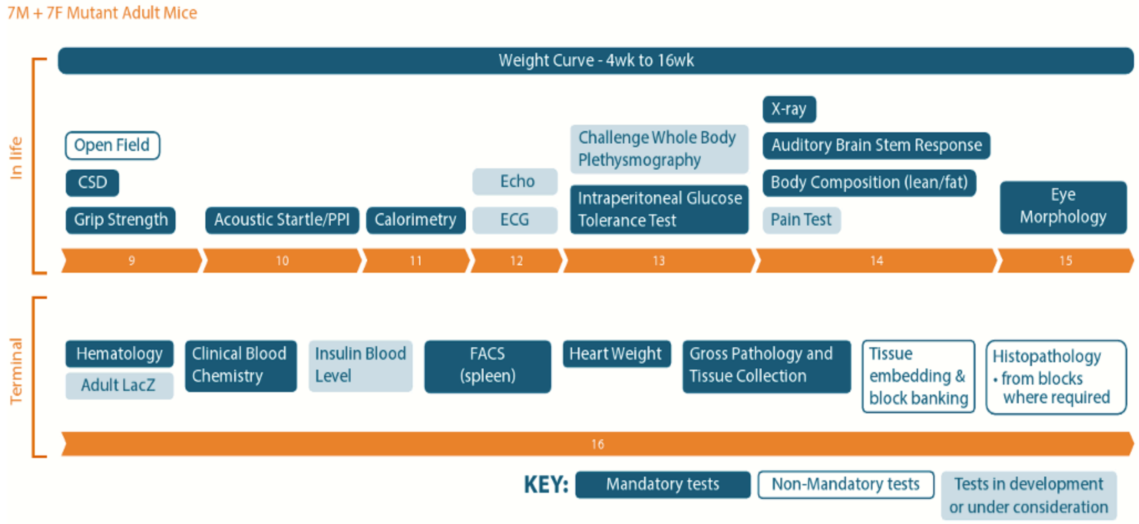


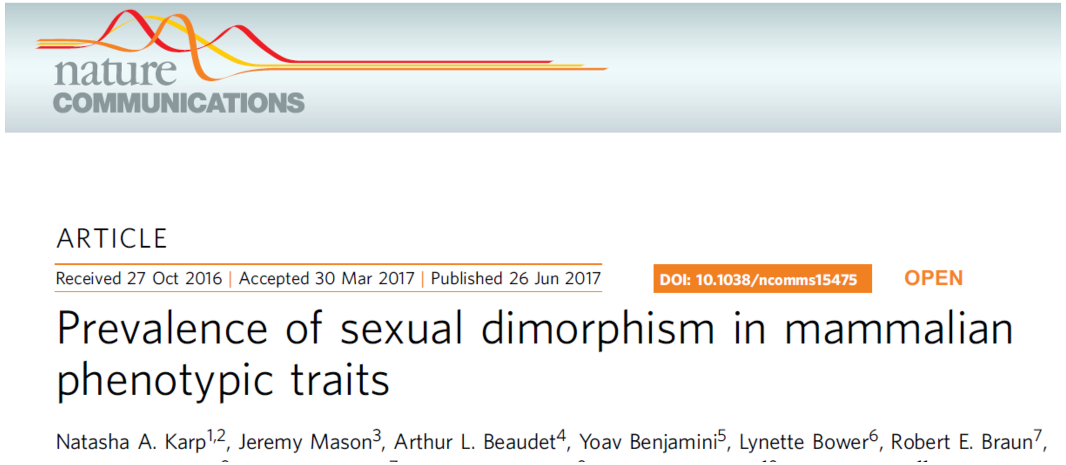

nature COMMUNICATIONS

ARTICLE

Received 27 Oct 2016 | Accepted 30 Mar 2017 | Published 26 Jun 2017

DOI: 10.1038/ncomms15475 OPEN


#### Prevalence of sexual dimorphism in mammalian phenotypic traits


Natasha A. Karp[1,2], Jeremy Mason[3], Arthur L. Beaudet[4], Yoav Benjamini[5], Lynette Bower[6], Robert E. Braun[7],


- 10 institutes
- 14,250 wildtype mice
- 40,192 mutant mice
- 2186 mutant lines
- up to 234 traits.

### SABV?

#### In control data

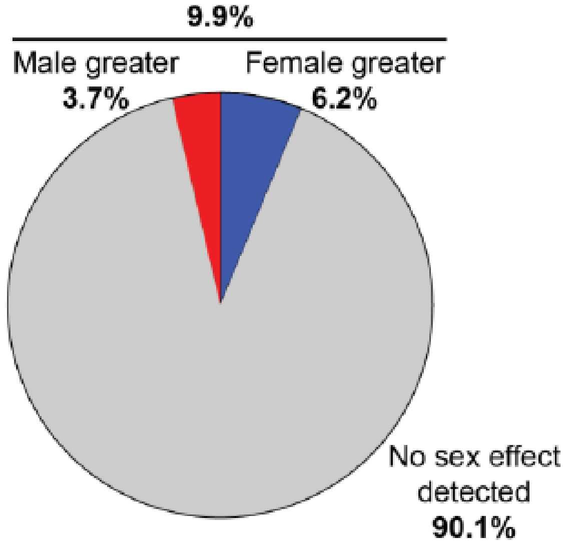


Categorical
N=545

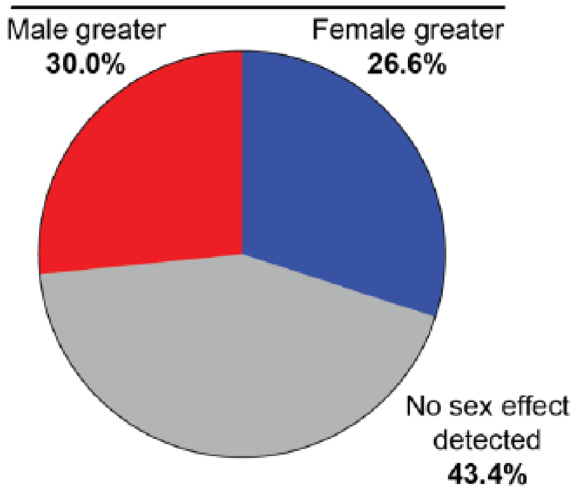


Continuous
N=903

#### As a modifier of treatment effect?

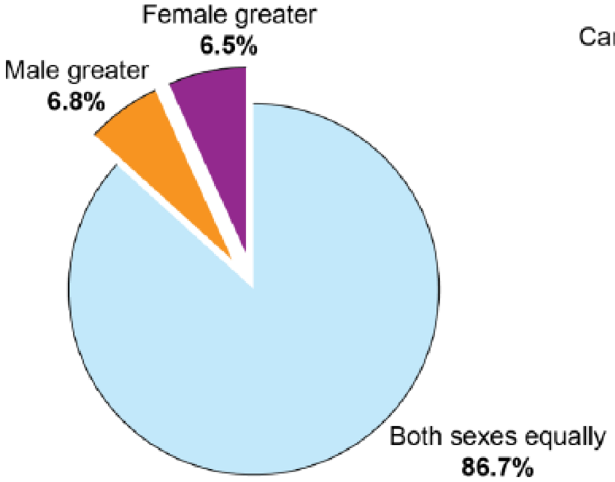


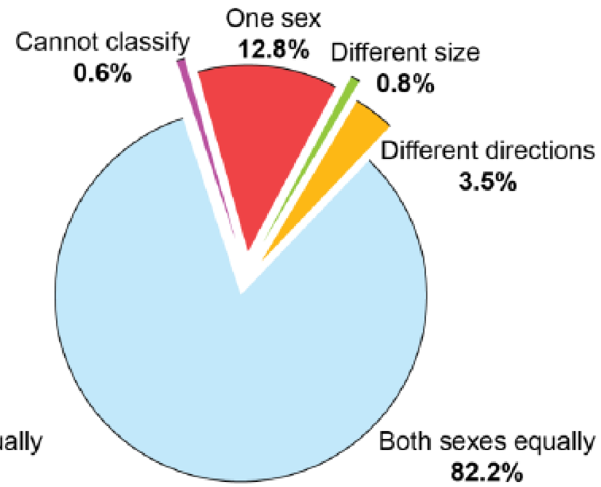


Categorical

No ds = 266,952
No ds sig = 1,220

Continuous

No ds = 110, 586
No ds sig = 7 929

*Karp et al 2017 Nature Communications*

### Sociological exploration of the issue

- Interviews to probe scientists’ (n=9) thoughts and experiences

**Generalizability**

- Important to embrace variation to understand biological differences

**Avoiding complexity**

- To make progress in science reduce complexity

**Practicality**

- Mediate tension between generalizability and avoiding complexity
- Convert the original research question into a doable problem
- Pragmatic constraints of which variables can be considered
- Availability of materials and cost constrained ability to include SABV

Gompers, Annika. Genderscilab, 2018.

### MRC survey

- Majority of MRC researchers (95% of in vivo researchers and 88% of cell users) saw benefit of considering diversity
  - Translatability
  - Reproducibility
  - Detecting sex specific effects

  - Barriers/Concerns
    - Cost of experiments and complexity of research design
    - Compliance with the principles of the 3Rs (animal usage for *in vivo* researchers)
    - Commercial availability of samples (cell researchers)

https://www.ukri.org/wp-content/uploads/2022/03/MRC-090322-SexInExperimentalDesign-SummaryReport.pdf

4

### What is sex inclusive research?

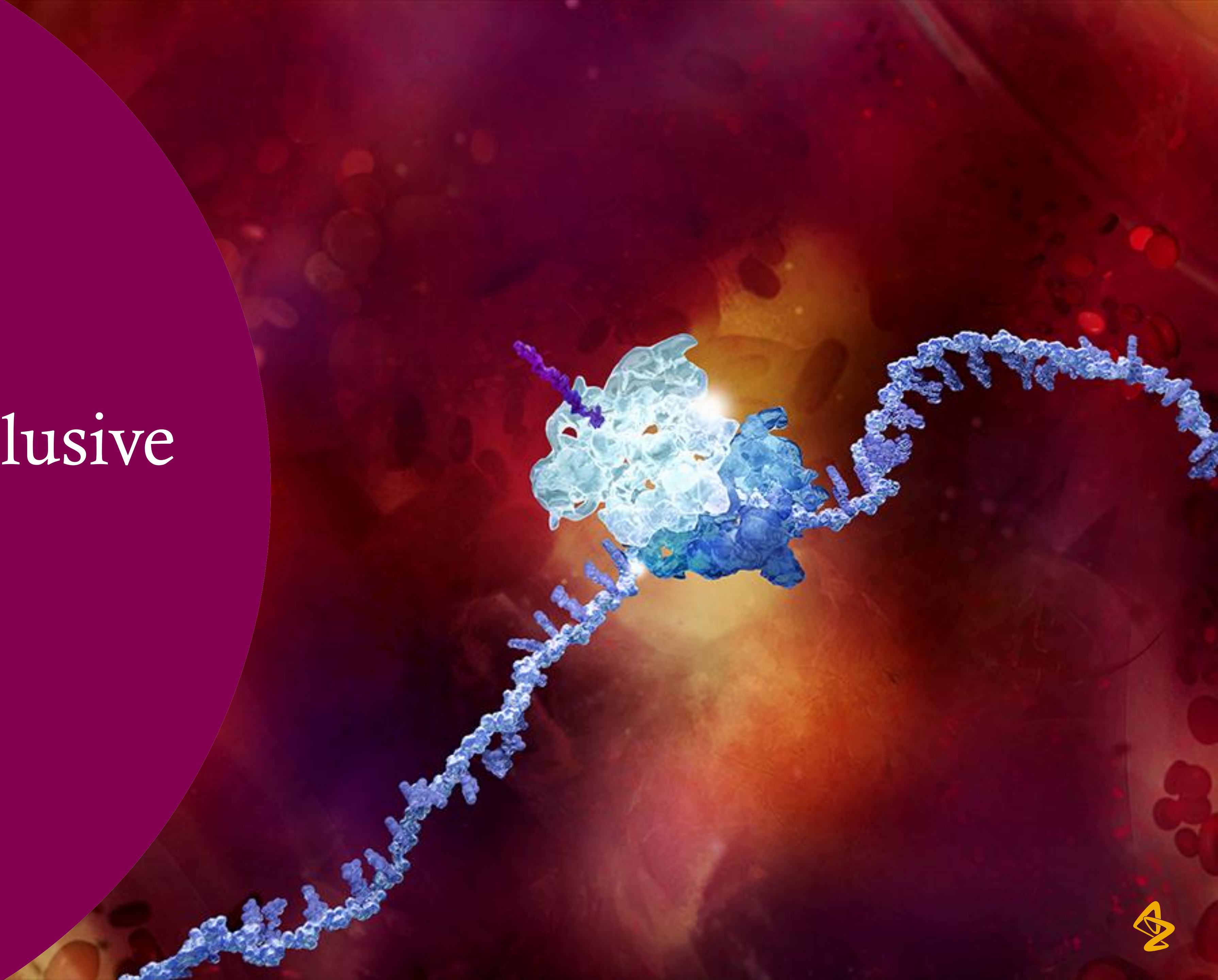

### Sex and Gender Equity in Research (SAGER) guidelines

- Principles
  - Use the term sex and gender carefully
  - Where the subjects of research comprise organisms capable of differentiation by sex, the research should be designed and conducted in a way that can reveal sex-related differences in the results, even if these were not initially expected.
  - Where subjects can also be differentiated by gender (shaped by social and cultural circumstances), the research should be conducted similarly at this additional level of distinction

Heidari 2016 Research Integrity and Peer Review

### Funding bodies are driving change

- Movement from recommendation to requirement and active questions in funding process

| Body | Year |
|---|---|
| NIH | 2016 – required incorporation both *in vivo* and *in vitro* |
| Canadian Institute of Health Research | 2010 – questions in grant application |
| Irish Research Council | 2013 – questions in grant application |
| European Commission | 2020 – required incorporation both *in vivo* and *in vitro* |
| MRC | 2022 – inclusion of both sexes the default for *in vivo* and *in vitro* |
| CRUK | *2023 – inclusion of both sexes the default for animal, tissues or cells* |

- WT funding - MESSAGE (Medical Science Sex and Gender Equity) project
  - Co-develop a sex and gender policy framework for funders and regulators in the UK

### Inclusion is the default – exclusion by exception

**Requirement**

- Specify the sex for human or animal tissues and cells used in experiments
- Inclusion of both sexes as default for studies involving animals and human and animal tissues and cells
- Analysis should account for sex

**Vision**

- Experiments are powered to detect the effect of interest across the two sexes
- Research will then estimate a **generalisable** result
- If the effect is very different between the sexes then this will become apparent

### Exception?

- Where sex cannot be determined
- Pure molecular studies such as P-P interactions
- Sex-specific conditions or phenomena e.g. ovarian cancer
- Acutely scare resources (e.g.  rare disease)
- If you can provide strong justification.

5

### Factorial analysis is the critical element

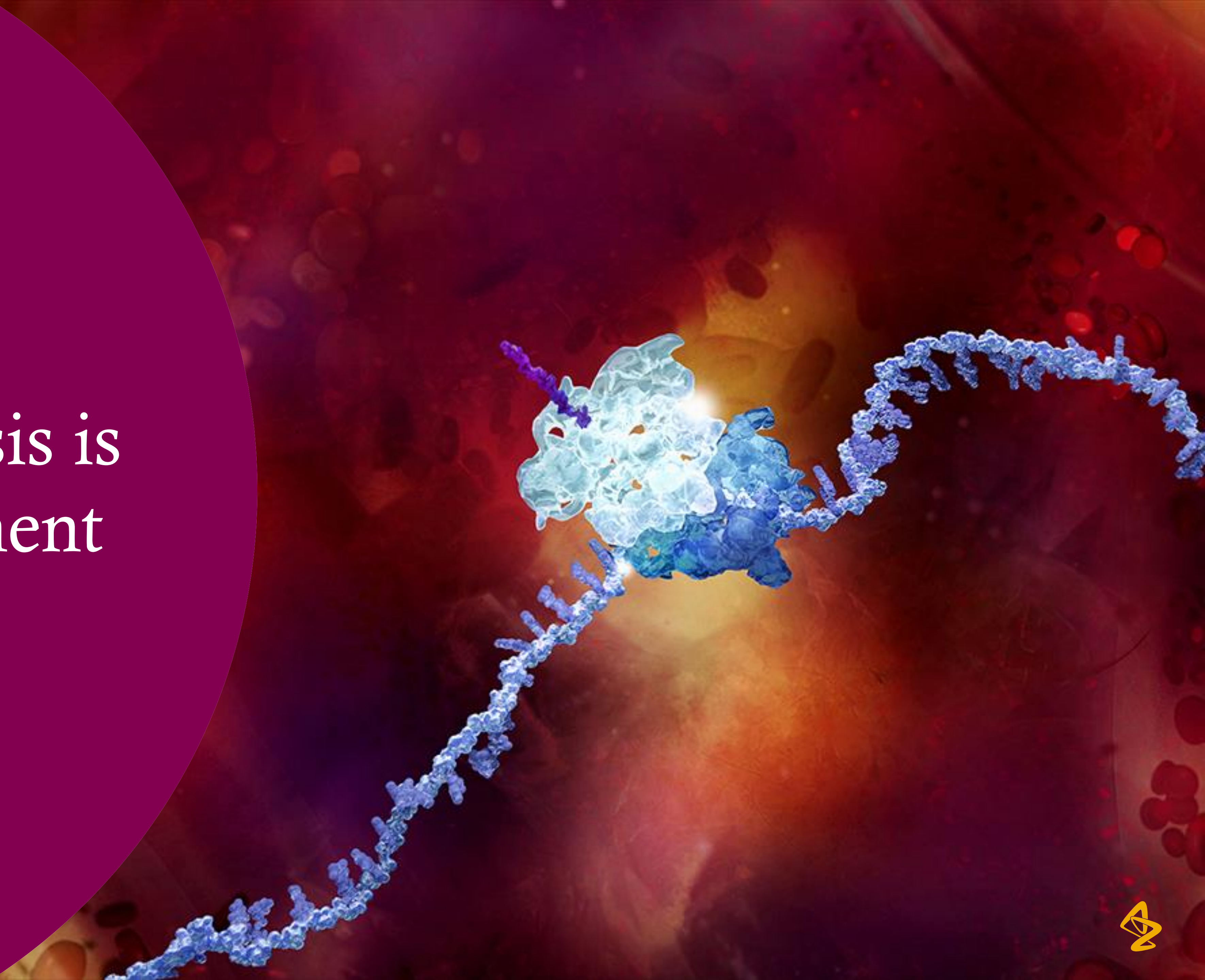

### Statistics as understanding sources of variation

- Some of this variation is background/impractical to explain. Some of it is planned (e.g., treatments) and systematic.
- We often aim to explain variability in the data through predictors that we control or observe:

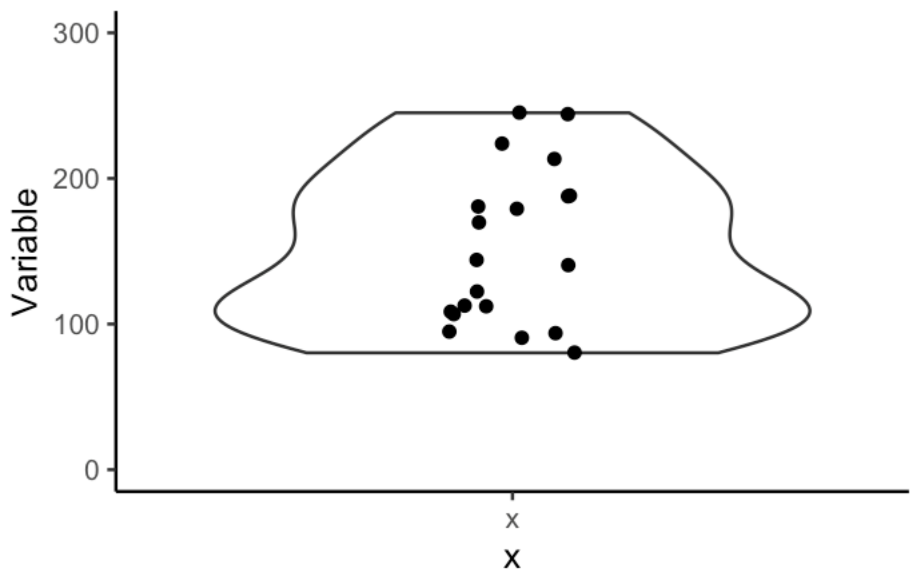


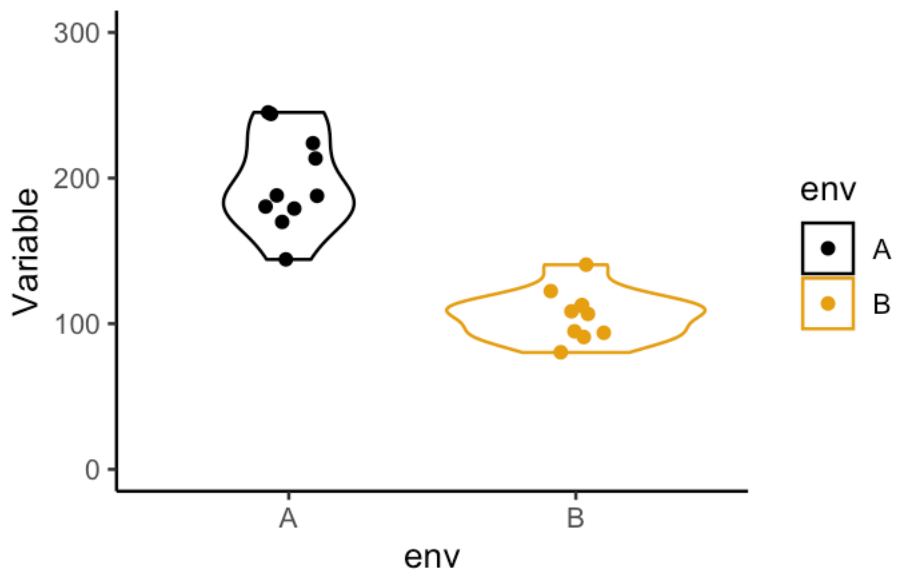

### Common sources of variation

#### Experiment

- Treatment
- Dose level
- Time
- Shelf life
- Instrument calibration
- Sampling

#### Environment

- Temperature
- Noise
- Light
- Smell
- Cage Size
- Bedding
- Enrichment activity
- No. of animals
- Diet
- Handling

#### Animal

- Species
- Strain
- Sex
- Batch
- Age
- Weight
- Health status

#### People

- Technician
- Surgeon
- Researcher

### Managing sources of variation

**Known sources:**

- Control (i.e. standardise)
- Block
- Factor of interest

**Unknown sources:**

- Randomize

### Statistical strategies- t-test vs block vs factor

#### T-test

Simplest approach: compare two conditions

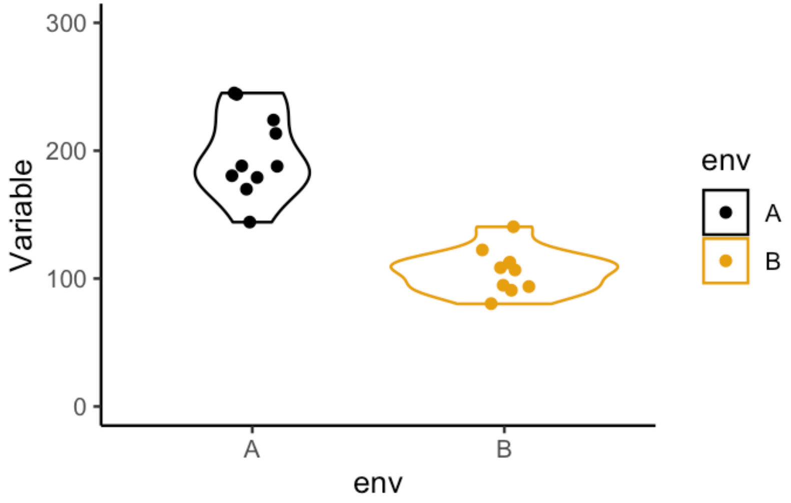


Y ~ environment

#### Block

When the interaction is not of biological interest:

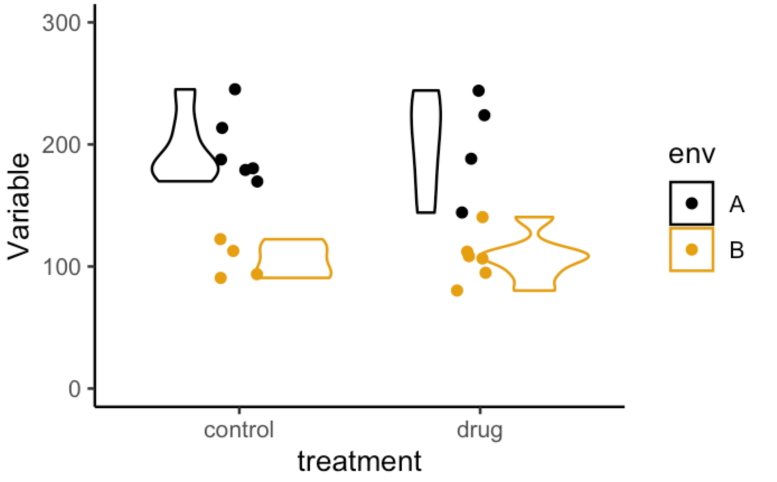


Y ~ treatment + environment

#### Factor

When the differential effect is biologically interesting:

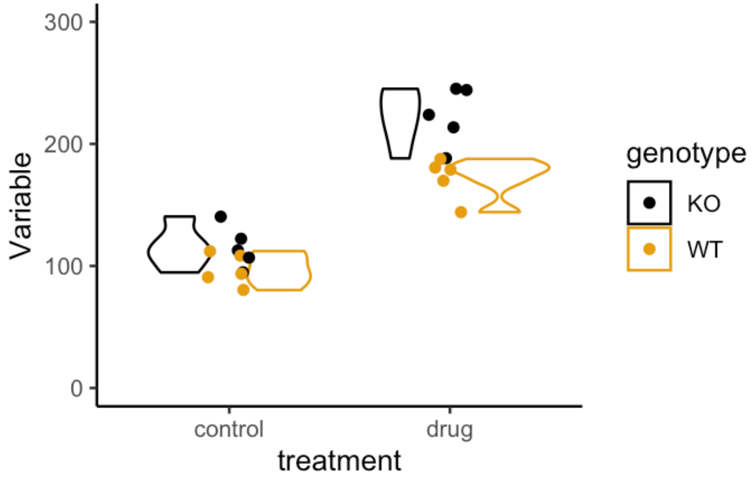


Y ~ treatment + genotype + treatment:genotype

### Moving from complete randomised to factorial design

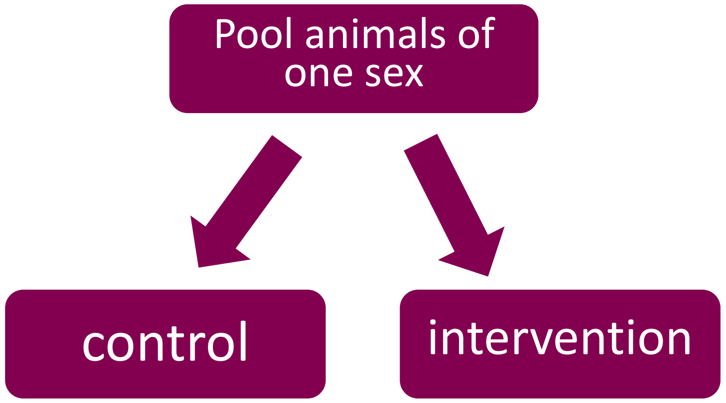


Y ~ treatment

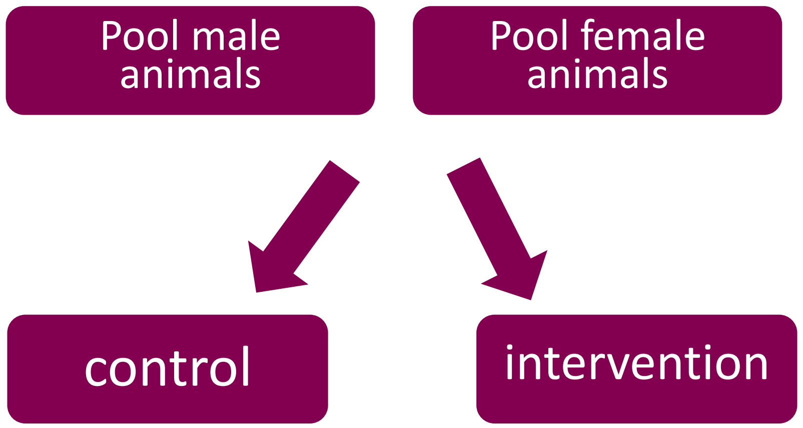


Y ~ treatment + sex + treatment*sex

### Testing the main effects: treatment and sex

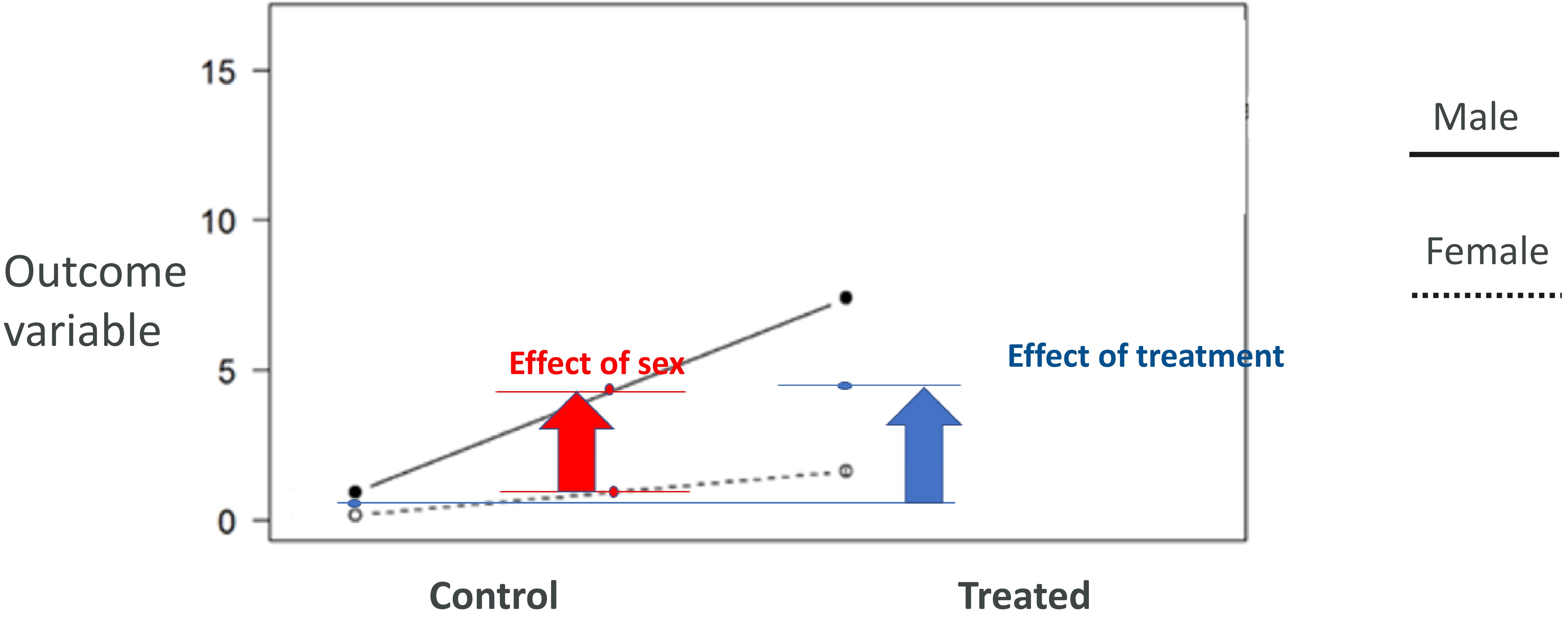

### Testing the interaction: comparison of differences

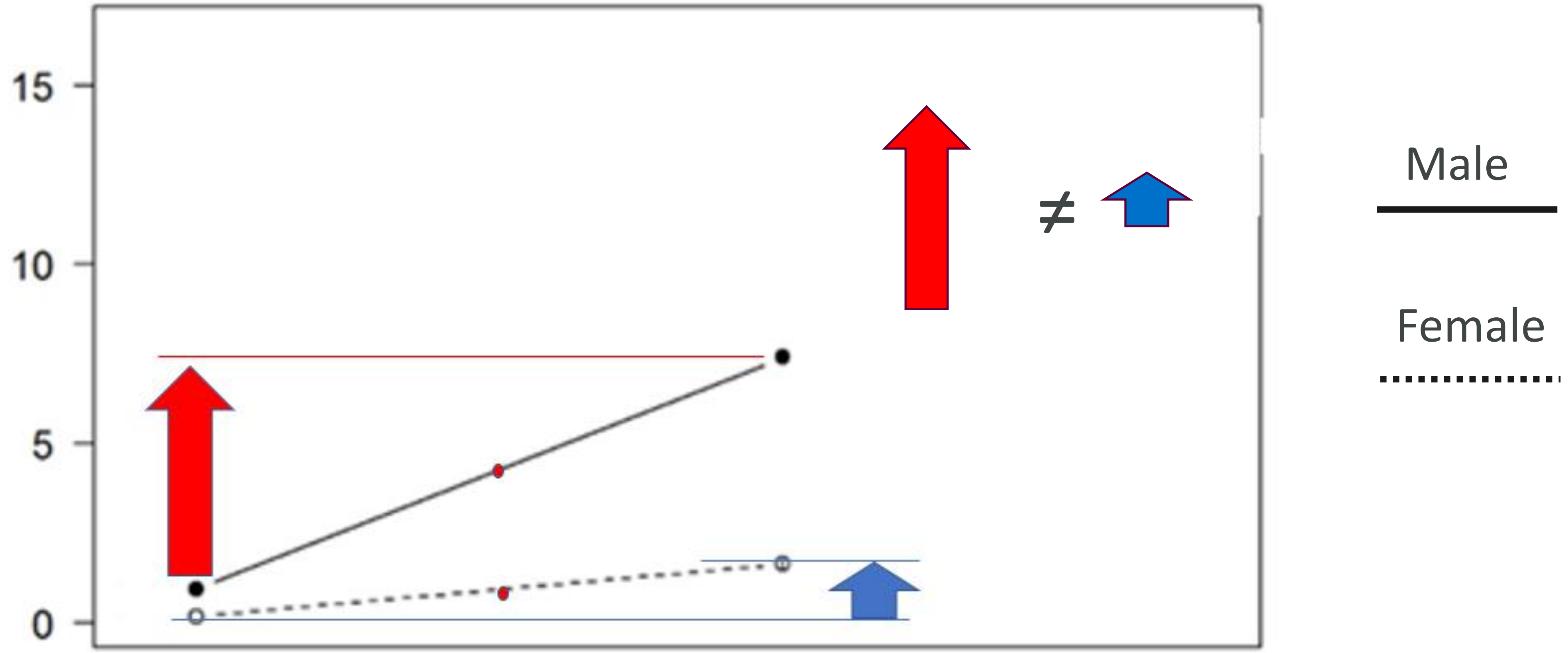


Outcome variable

**Control** **Treated**

### Some examples

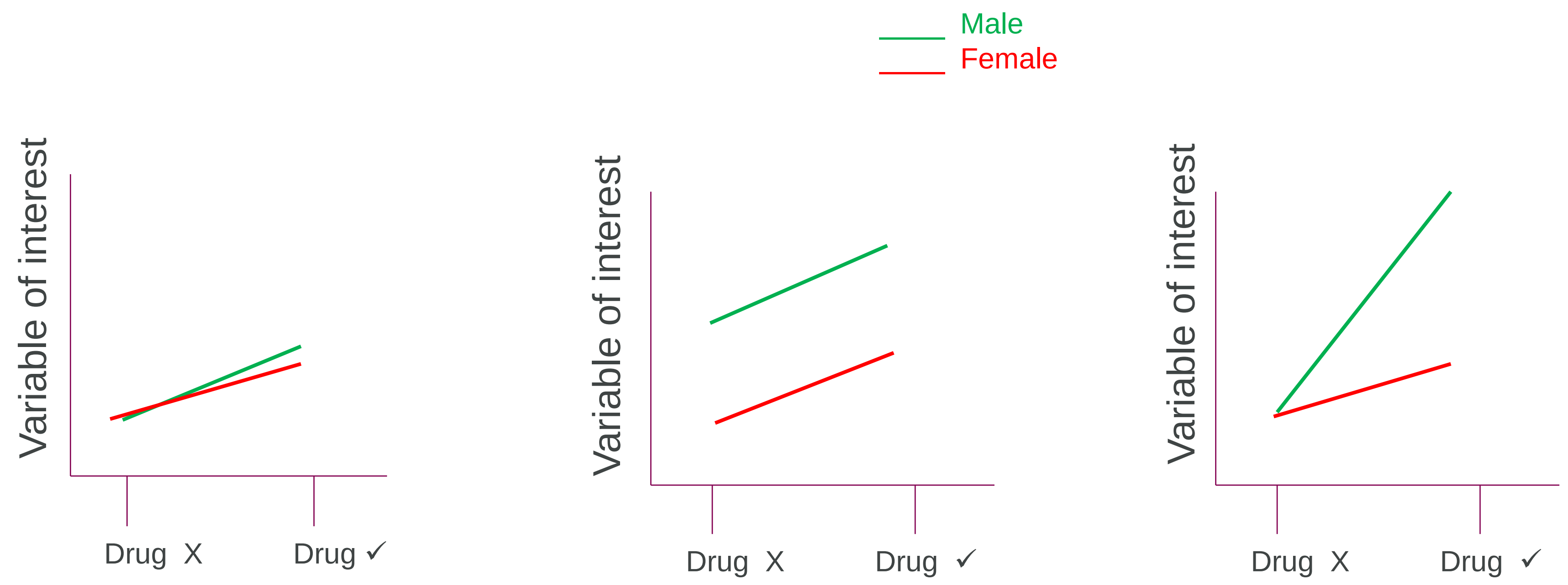


| Effect | Significant |
|---|---|
| Sex | X |
| Drug | ✓ |
| Interaction | X |

| Effect | Significant |
|---|---|
| Sex | ✓ |
| Drug | ✓ |
| Interaction | X |

| Effect | Significant |
|---|---|
| Sex | ✓ |
| Drug | ✓ |
| Interaction | ✓ |

### Misconception: It will increase my animal usage

"Keep doing what you are already doing but change half the animals in your study to female"

*McCarthy **2015** Schizophrenia Bulletin*

- **Evidence:**
- **Including both sexes in *in vivo* research does not necessitate an increase in sample size: a key role for factorial analysis methods** (Phillips PLoS Biology 2023)
- **Inclusion of females does not increase variability in rodent research studies** (Beery Curr Opin Behav Science 2018)

### The interaction term is critical

"We will test the females once we understand the biology"

- Can't tell if the effect size is significantly different
- You never get to the second sex
- It is a structurally biased research approach

  - "false assumption that what is discovered in males is how the brain really works, whereas in females, the same neurobiological processes are probably more complicated. "
  - "When females are studied through a male lens, the true crux of the research question for females can be missed. "
    - ARE HORMONES A "FEMALE PROBLEM" FOR ANIMAL RESEARCH?
    - https://www.science.org/doi/10.1126/science.aaw7570

### Common mistakes & how to avoid them – misidentification of design (1)

It can be easy to misidentify the study design and use the wrong analysis as a result. These mistakes are common when researchers report studies involving both sexes:

**Pooling**

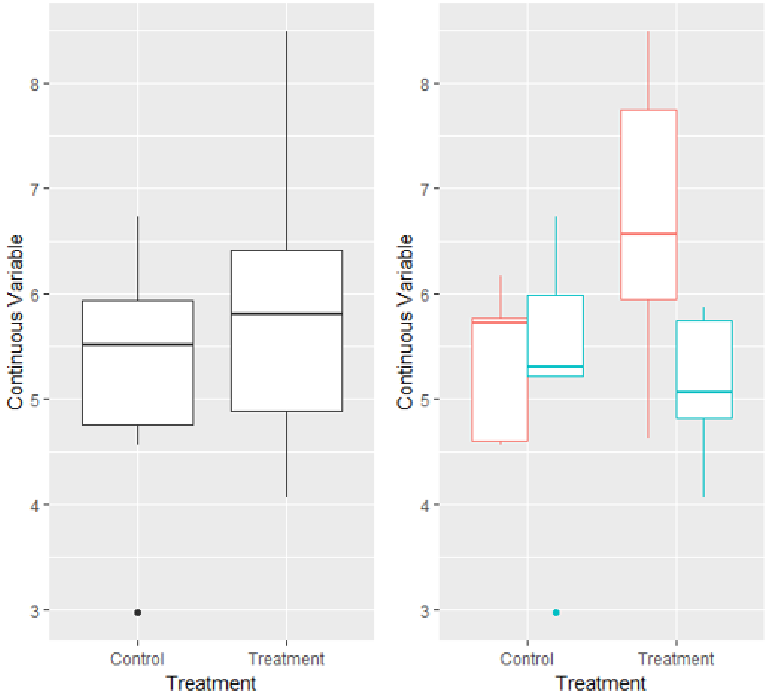


**Disaggregation**

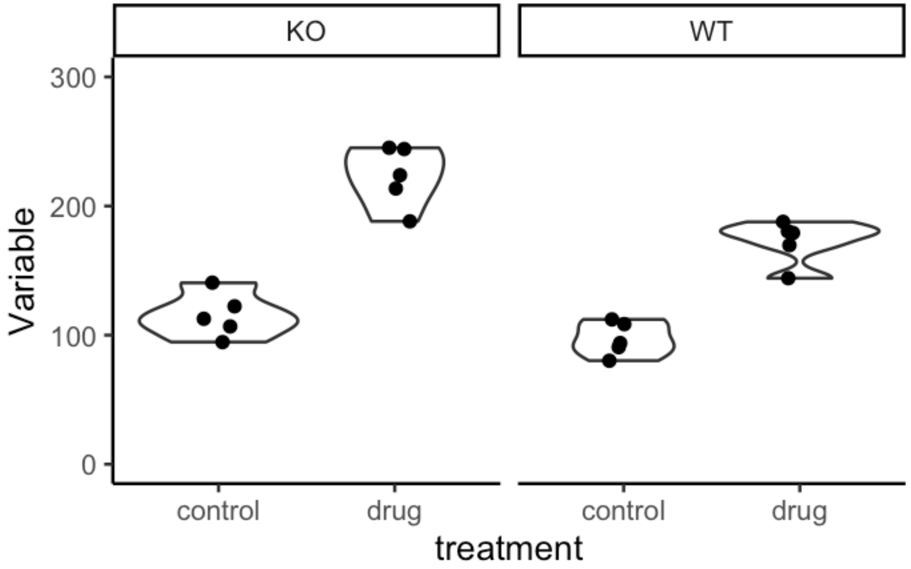


*Garcia-Sifuentes & Maney, eLife 2021*

### Common mistakes & how to avoid them- misidentification of design (2)

Another commonly seen error- treating a factorial design as a parallel group design and using a one-way ANOVA: Genotype * Treatment design:

#### Mistake:

- Fails to identify experimental structure- loss of power & fails to test the biology of interest:

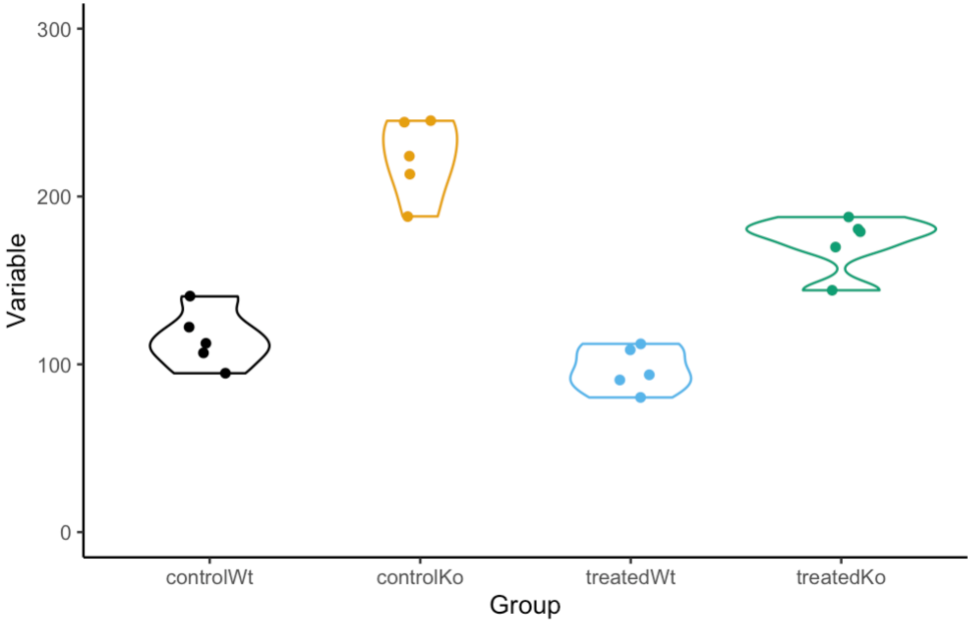


#### Correct:

- Enables estimation of interaction and main effects:

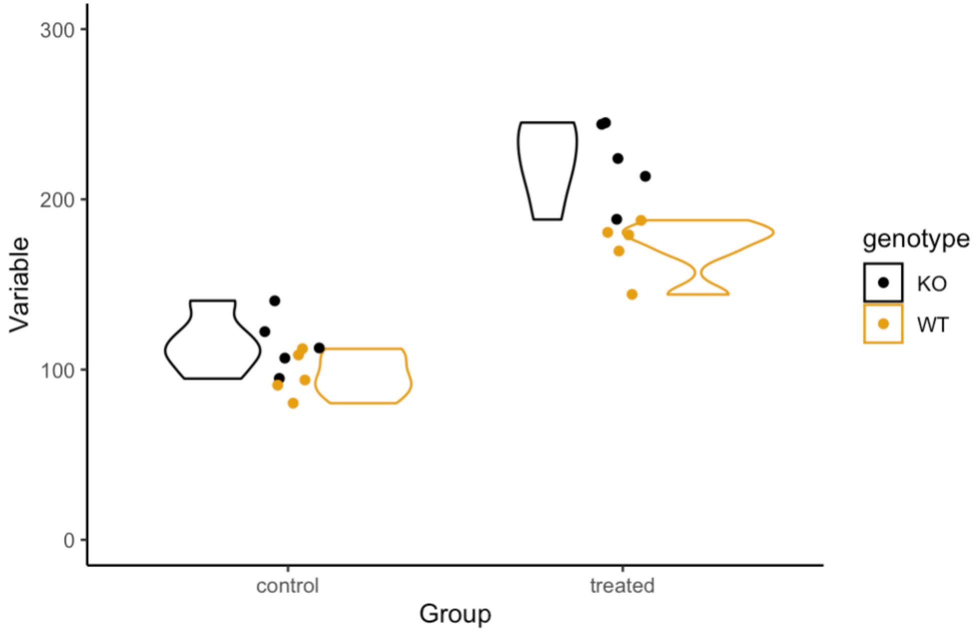

### A simulation-based approach for exploring factorial power

1) Generate simulated data
2) Apply statistical test and extract p values
3) Repeat process 1000 times for each effect size of interest and calculate the % of significant p values as the statistical power for that scenario.

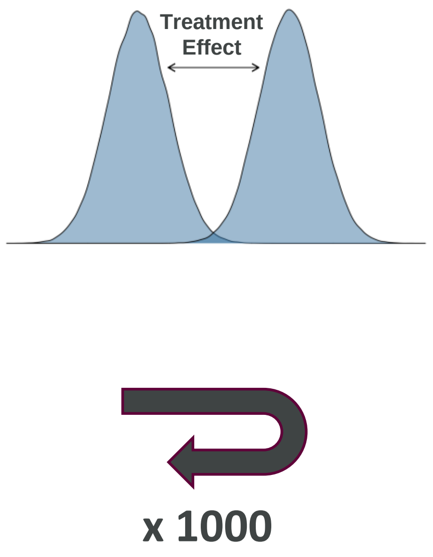


**Scenarios**

1. Baseline effect of sex
2. A differential effect of sex on treatment in the same direction
3. An effect in one sex only
4. Opposite effects of sex by treatment

Phillips 2023 PLOS Biology

### 1: When there is a baseline effect of sex?

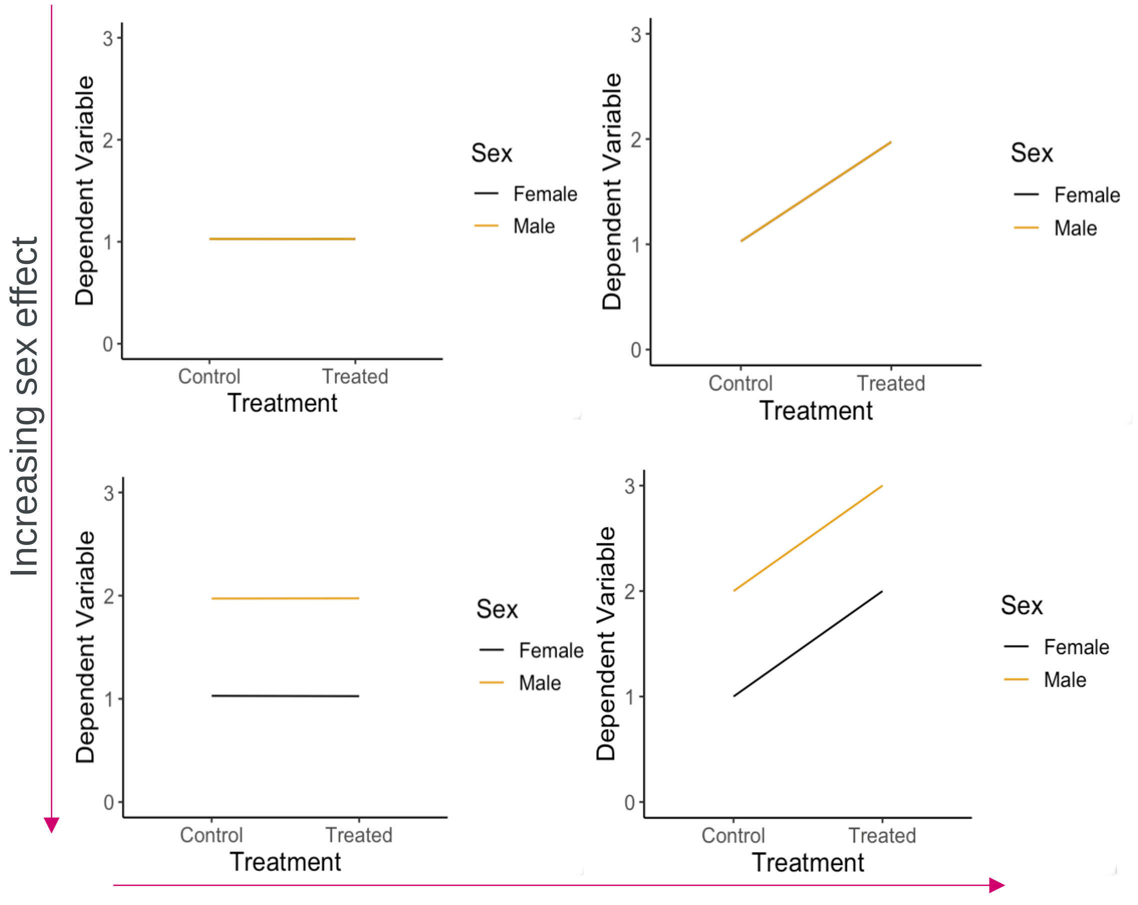


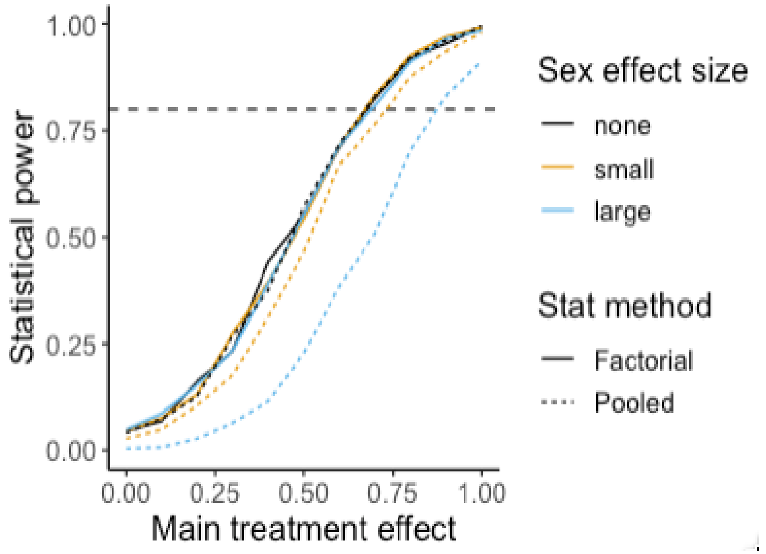


Phillips 2023 PLOS Biology

### 2: When there is a differential effect in the same direction?

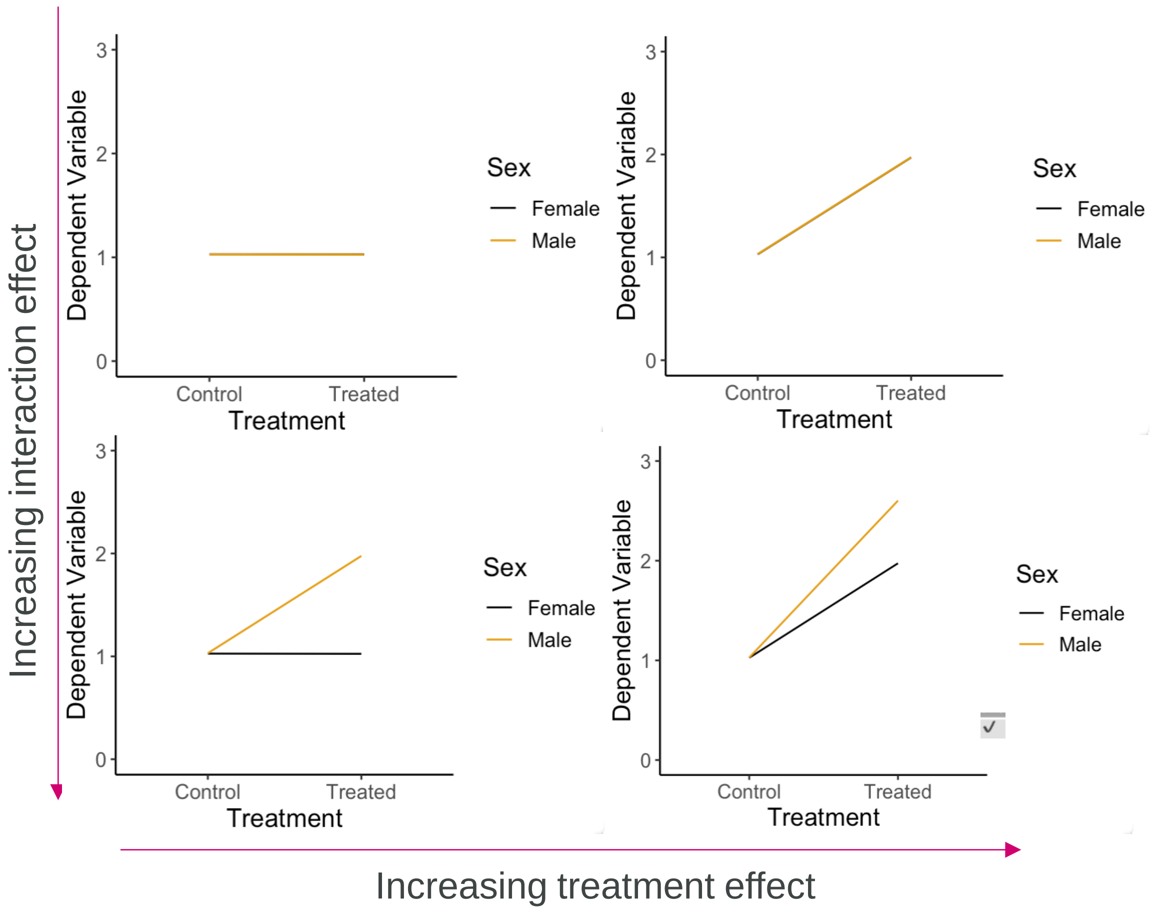


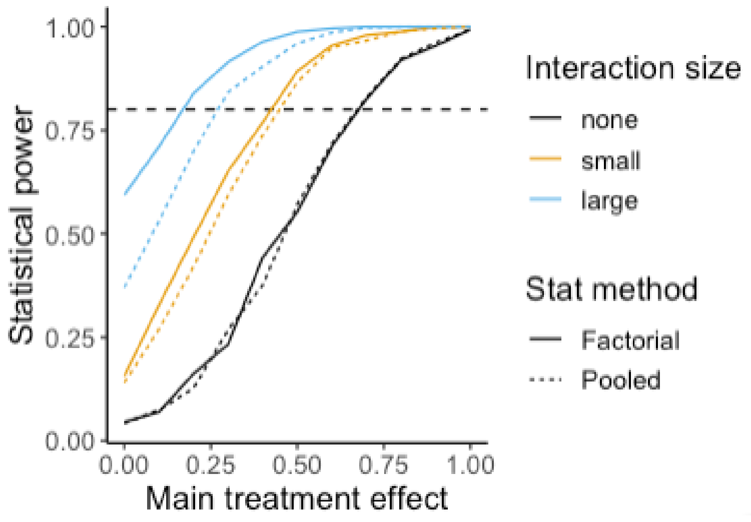


Phillips 2023 PLOS Biology

### Pooling compromises treatment power when there is an effect in one sex

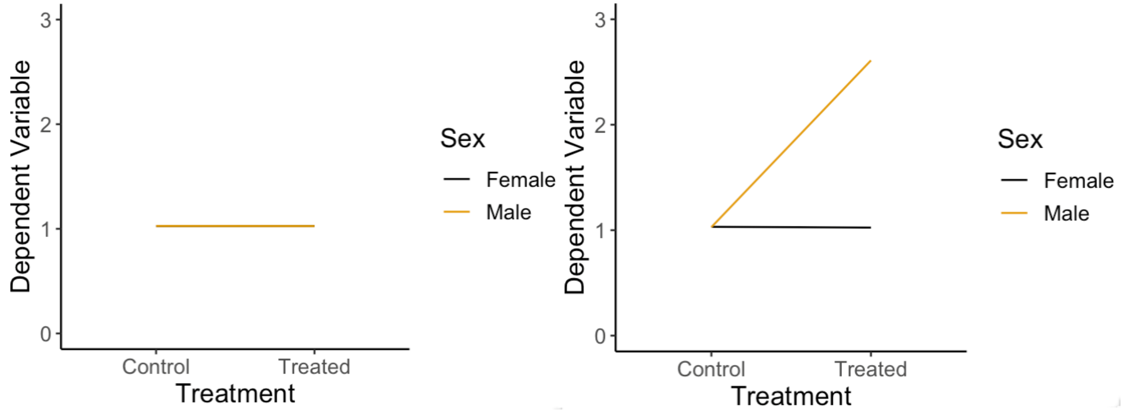


Increasing interaction effect (males only)

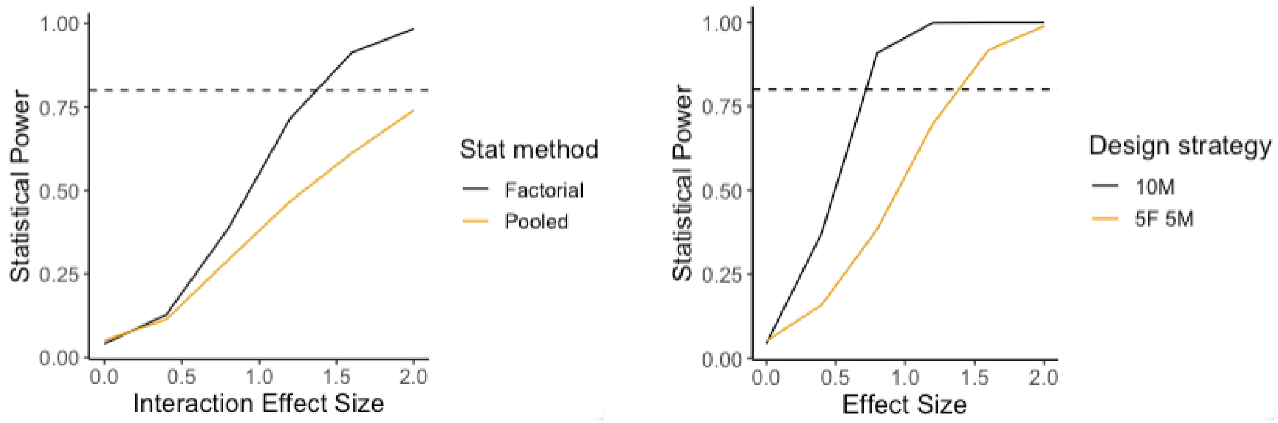


Phillips 2023 PLOS Biology

### Power is passed to the interaction term when the effects are in opposite directions

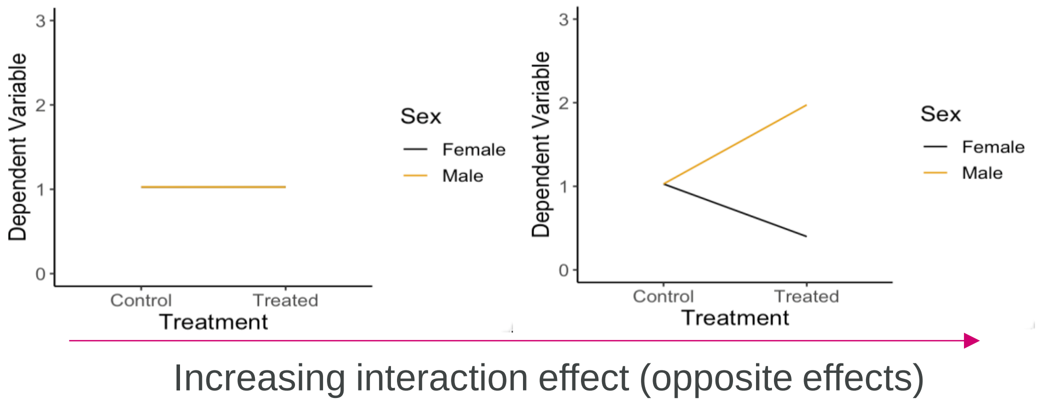


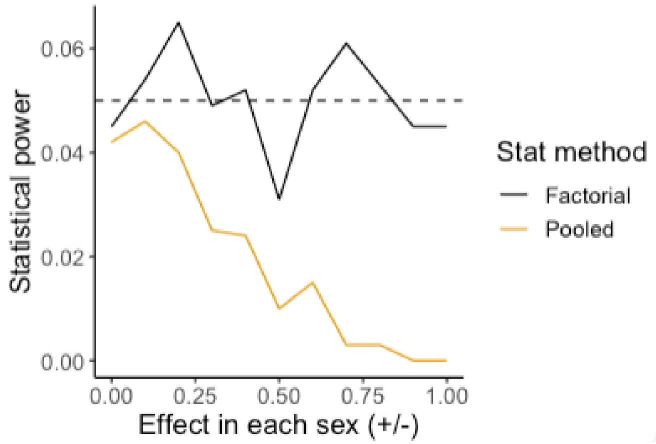


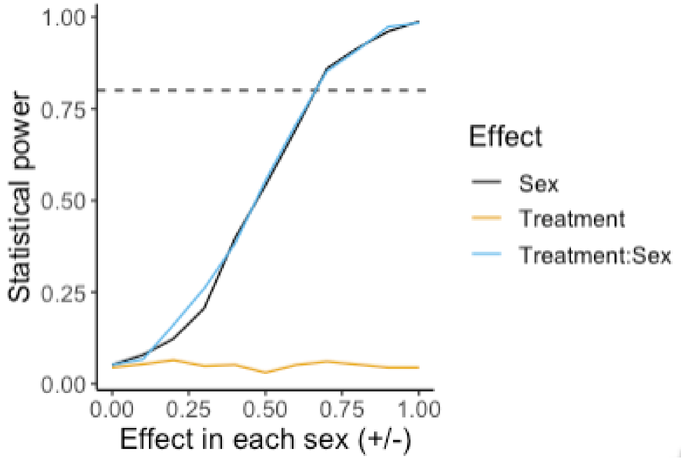


Phillips 2023 PLOS Biology

### Simulation conclusions:

A factorial approach provides power benefits, compared to a pooling t-test approach.

Disaggregation of data by sex loses statistical power and doesn't test for the interaction.

Typically, there is no loss of power to detect a treatment effect when including both sexes.

When power is lost, the knowledge gained is vital as the power is transferred to the interaction term.

### Interactive activities

### Question 1

You are working on understanding the effect of Compound X on blood pressure in rats. To improve the generalizability of the study, you have decided to test a vehicle control vs Compound X in 3 different inbred strains of rat. You are not interested in whether the effect of Compound X differs across strains.

Which analysis strategy would you select for this design?

1. Pooling the data for an intervention and then running a t-test between treated and control data.
2. Disaggregating the data and completing separate t-tests between treated and control data for each strain.
3. A two-way ANOVA testing for main effects only.
4. A two-way ANOVA testing for main effects and an interaction.

### Question 2

You are working on understanding the effect of Compound X on blood pressure in rats. Based on the literature, you suspect that the drug may have a greater effect in females. You have decided to test vehicle control vs Compound X in both male and female rats.

Which analysis strategy would you select for this design?

1. Pooling the data for an intervention and then running a t-test between treated and control data.
2. Disaggregating the data and completing two separate t-tests between treated and control data.
3. A two-way ANOVA testing for main effects only.
4. A two-way ANOVA testing for main effects and an interaction.

### Question 3

You have run an experiment in both male and female mice, testing the effect of a drug designed to improve cardiac disease. You have obtained the following results.

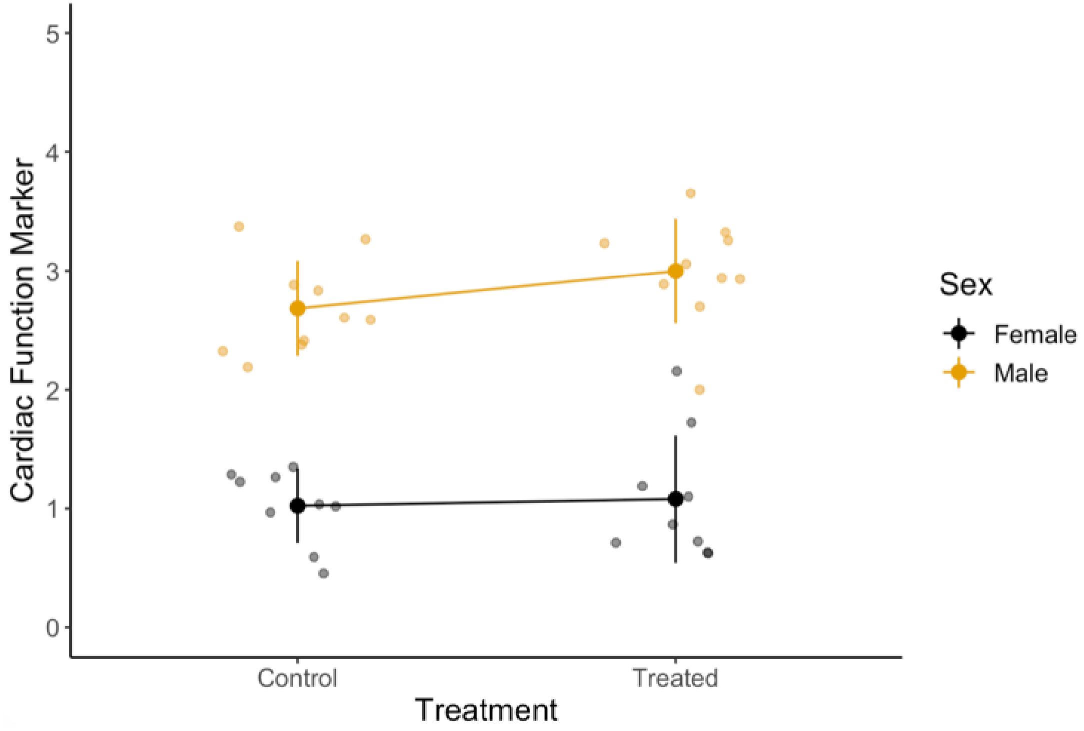


1. From these data, predict the results obtained from each term of a two-way ANOVA (sex, treatment, interaction).
2. What is the biological conclusion?

### Question 4

4.You have run an experiment in both male and female mice, testing the effect of a drug designed to improve cardiac disease. You have obtained the following results.

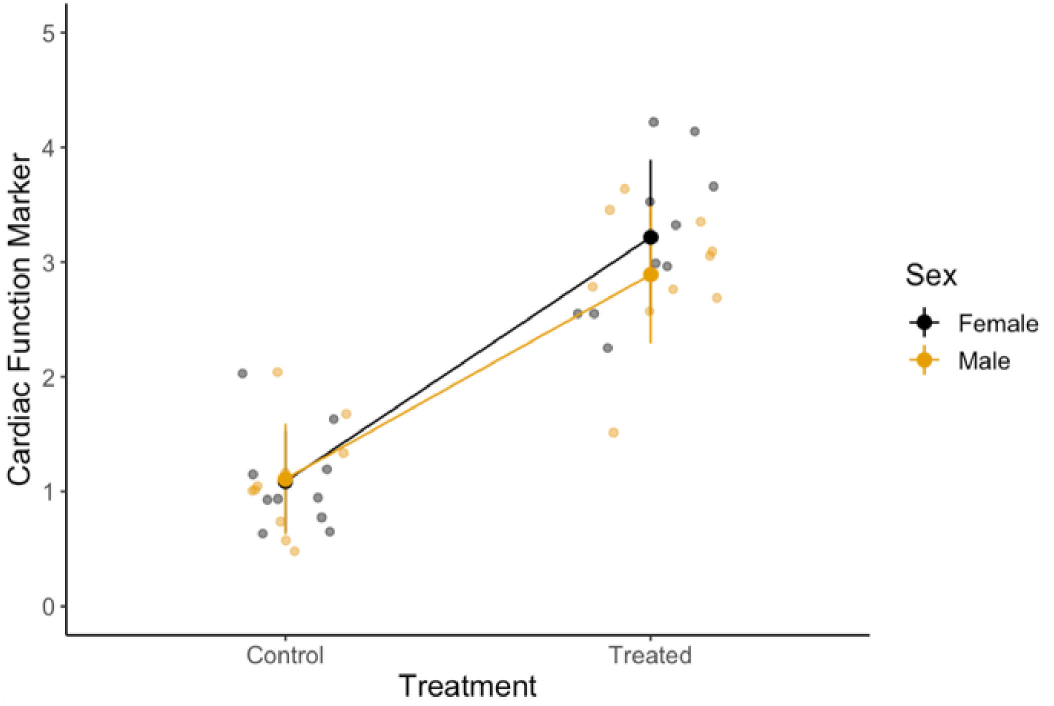


1. From these data, predict the results obtained from each term of a two-way ANOVA (sex, treatment, interaction).
2. What is the biological conclusion?

### Question 5

You are testing the effect of knocking out a certain gene on a marker of inflammation in male and female mice. You have obtained the following results. The interaction term between sex and genetic status is statistically significant.

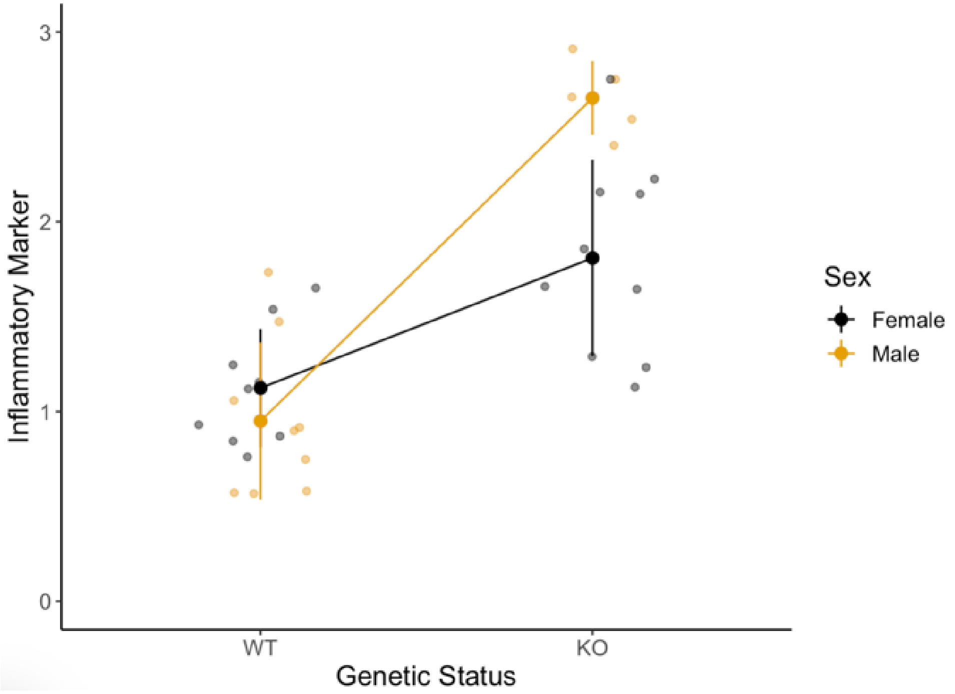


1. From these data, predict the results obtained from each term of a two-way ANOVA (sex, treatment, interaction).
2. What is the biological conclusion?

### Question 6:

Scenario: We have run an experiment where there is a baseline sex difference and the intervention increases the outcome variable. Which statistical method will be more sensitive to detect the intervention effect?

1. Pooling the data: A single t-test between treated and control.
2. Disaggregating the data by sex and a separate t-test between treated and control
3. A two-way ANOVA testing for main effects and an interaction.

6

British Pharmacological Society

BJP British Journal of Pharmacology

REVIEW ARTICLE | Free Access

**Sex bias in preclinical research and an exploration of how to change the status quo**

Natasha A Karp, Neil Reavey

First published: 12 November 2018 | https://doi.org/10.1111/bph.14539 | Citations: 20

This article is part of a themed section on The Importance of Sex Differences in Pharmacology Research. To view the other articles in this setion visit http://onlinelibrary.wiley.com/doi/10.1111/bph.v176.21/issuetoc

### Exploring the barriers

- MISCONCEPTIONS
- SKILL GAP
- PRACTICAL CONCERNS
- 3R INTERPRETATION OF REDUCTION

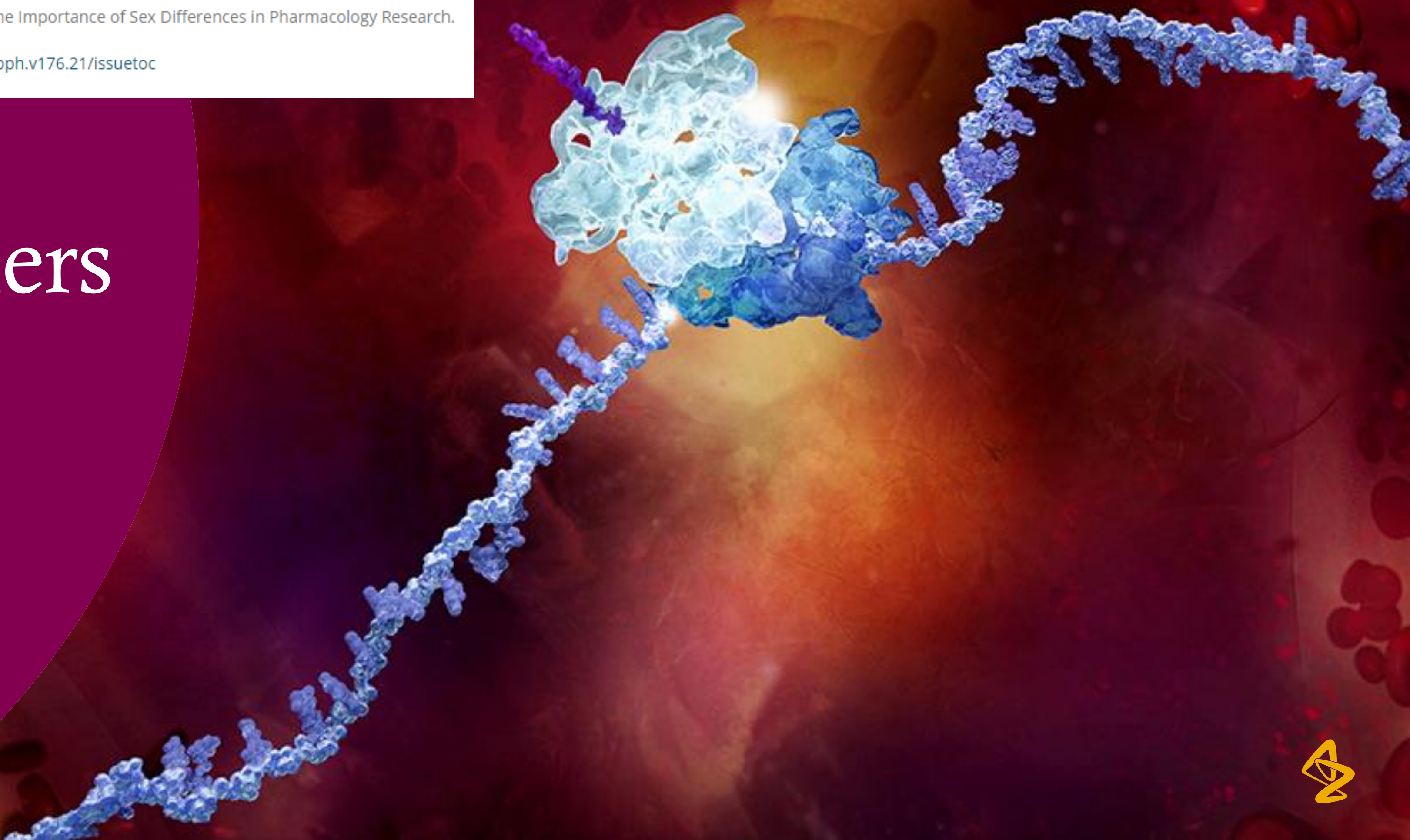

### Misconception: hormonal cycles: females more variable

**Rats** Becker 2016 BSD

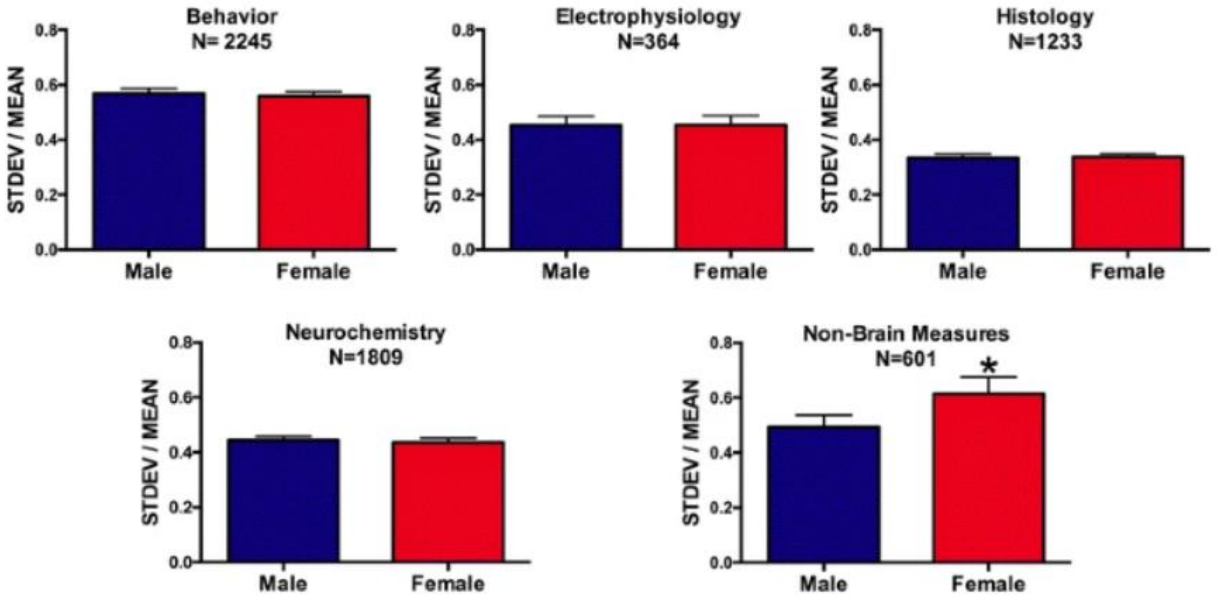


"Female rats were not more variable at any stage of the estrous cycle than male rats."

**Mice** Prendergast 2014 NNBR

- meta-analysis 293 published articles
- behavioral, physiological, morphological, and molecular traits
- CV distribution = no differences
- At trait level – for three types of traits males were more variable than females

"Randomly cycling female mice were no more variable than males on any trait."

### Inclusion isn't at odds with the 3R mindset

1. Breeding – produces both males and female animals

2. Reduction in N across experiments – more efficient to include both sexes

| | Standard | Contemporary |
|---|---|---|
| **Reduction** | Methods which minimise the number of animals used per experiment | Appropriately designed and analysed animal experiments that are robust and reproducible, and truly add to the knowledge base |

https://www.nc3rs.org.uk/the-3rs

### Fear of change

“To date, sex hasn’t explained variation in my model”

- Lack of data regarding sex differences does not indicate there are none
- The goal isn’t to identify sex differences but to estimate generalisable effects and be able to detect very large differences when they do occur
- Meta analysis has found that data analysis is often poorly conducted and hence historic conclusions can be misleading [https://elifesciences.org/articles/70817]

“My prior work has only considered in one sex”

- Unfortunately it carries lots of risk.
- “ To change is difficult.  Not to change is fatal”     William Pollard

My research to date has only studied one sex.

Why not both?

To include both would require the sample size to be doubled and that isn't ethical.

Research has shown that you can use a factorial design and you just share your N across both sexes and estimate the treatment effect from both

I can't do that because the effect would be different in the two sexes

Provided it is in the same direction, the factorial design would estimate the average effect and account for any baseline difference

No the effect is too different – I have to have an individual estimate. We also just don't know and it would be risky to assume it will the same

But when you are studying one sex you are assuming is the same. Is this the fear of the unknown and change in how we work? Opposite direction is biologically rare. Following the sex inclusive design would actually allow us to know when there is a big difference .

### Single sex justification

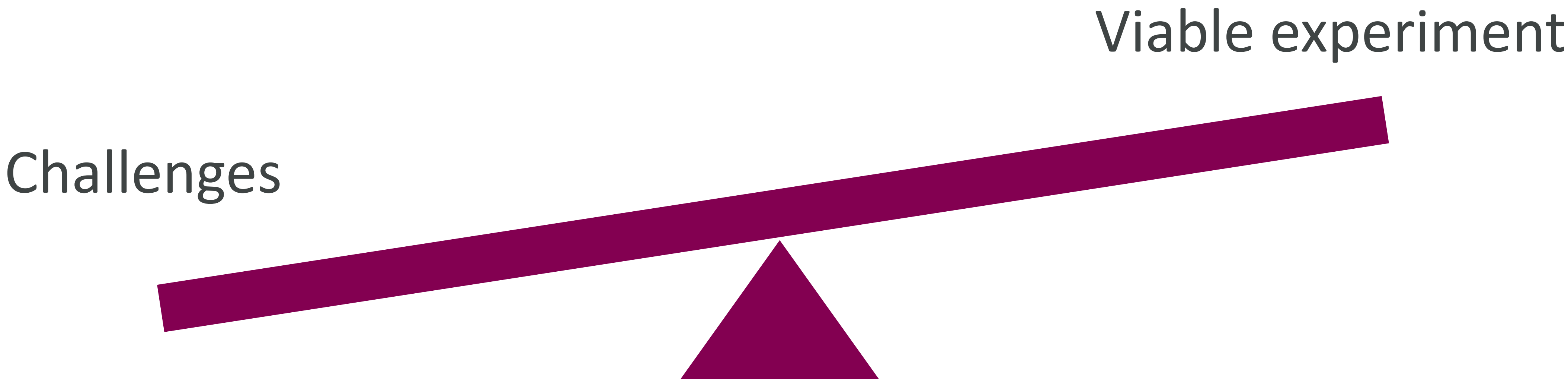


The justification could be appropriate following exploration for that study of logistical, ethical, or cost implications relative to the benefit of using both sexes of animals in a research proposal.

### SIRF: Sex Inclusive Research Framework

#### Why?

- Regulatory bodies assessing whether a research proposal is appropriate
- Frequently barriers are misconceptions
- Need transparency in the decision-making process
- We need educational resource to help move into considered justification to assess whether sex inclusion is a possibility.

#### What?

- Decision tree of ten questions and associated supporting information
- Delivers 1 or more classifications
- Options:
- **Green**: Proposal is appropriate
- **Amber**: Caution is required (I.e., the proposed design/analysis carries some risk)
- **Red**: Justification for single sex study in the proposal is not sufficient

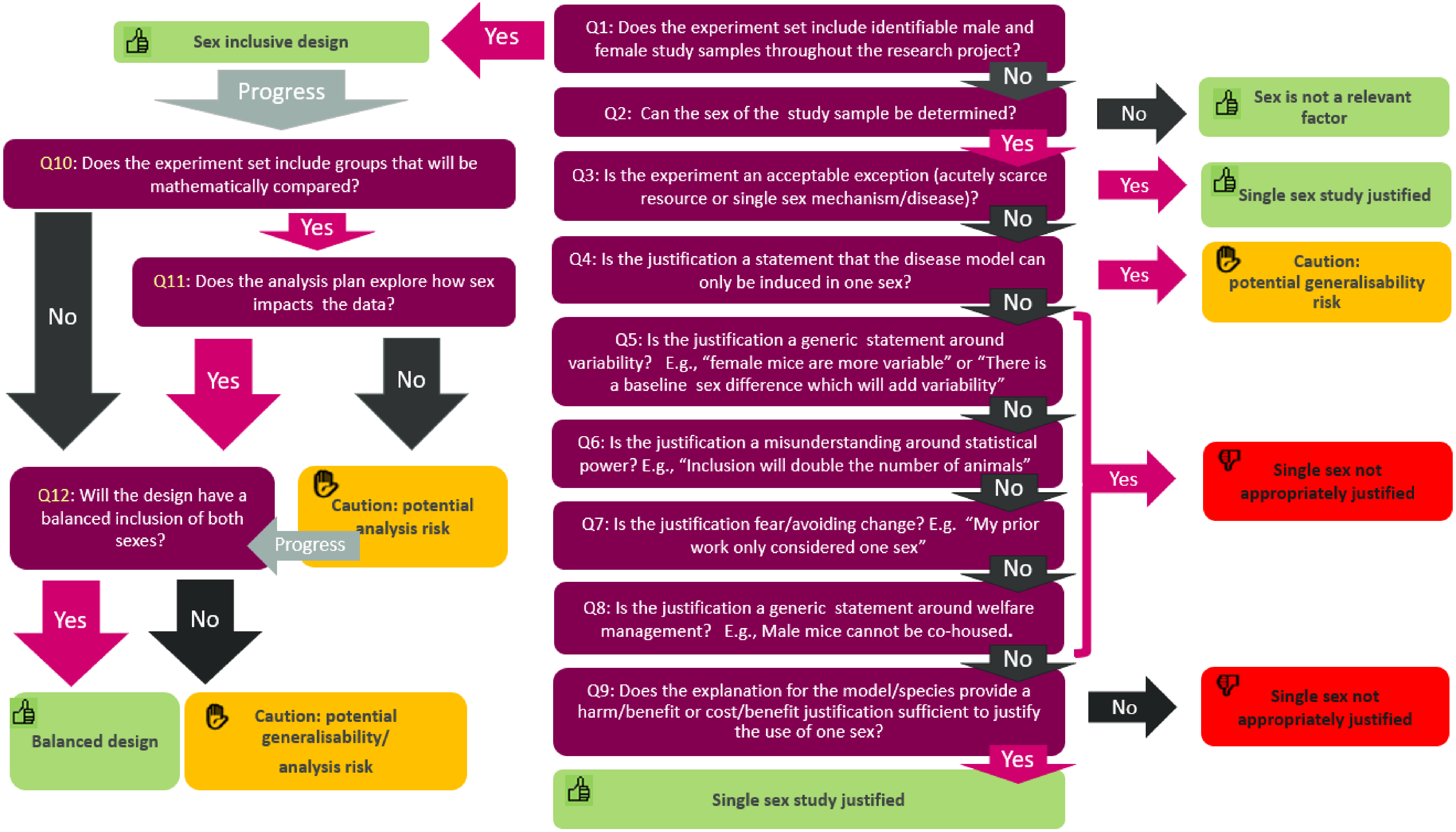

Q1: Does the experiment set include identifiable male and female study samples throughout the research project?
Yes
Sex inclusive design
Progress
No
Q2: Can the sex of the study sample be determined?
No
Sex is not a relevant factor
Yes
Q3: Is the experiment an acceptable exception (acutely scarce resource or single sex mechanism/disease)?
Yes
Single sex study justified
No
Q4: Is the justification a statement that the disease model can only be induced in one sex?
Yes
Caution: potential generalisability risk
No
Q5: Is the justification a generic statement around variability? E.g., “female mice are more variable” or “There is a baseline sex difference which will add variability”
No
Q6: Is the justification a misunderstanding around statistical power? E.g., “Inclusion will double the number of animals”
No
Q7: Is the justification fear/avoiding change? E.g. “My prior work only considered one sex”
No
Q8: Is the justification a generic statement around welfare management? E.g., Male mice cannot be co-housed.
Yes
Single sex not appropriately justified
No
Q9: Does the explanation for the model/species provide a harm/benefit or cost/benefit justification sufficient to justify the use of one sex?
No
Single sex not appropriately justified
Yes
Single sex study justified
Q10: Does the experiment set include groups that will be mathematically compared?
Yes
No
Q11: Does the analysis plan explore how sex impacts the data?
Yes
No
Caution: potential analysis risk
Progress
Q12: Will the design have a balanced inclusion of both sexes?
Yes
No
Balanced design
Caution: potential generalisability/ analysis risk

### Activity

- Use the framework to evaluate some published justifications for the design implemented [https://elifesciences.org/articles/56344]
- Determine whether the research was
  - **Green** – design was appropriate from a sex inclusive research perspective
  - **Amber** – research study raises some caution statements due to risk in the design/analysis
  - **Red** – design was not appropriate from a sex inclusive research perspective

### Slido voting!

- Participants can vote at Slido.com with **#7809431**

### Justification 1: we pooled the data because …..

"We could not compare the sexes as only four of the 25 wood mice were females. However, we have previously shown that the sex of wood mice has no effects on seed management "

Assessment options:

- **Green** – design was appropriate from a sex inclusive research perspective
- **Amber** – research study raises some caution statements due to risk in the design/analysis
- **Red** – design was not appropriate from a sex inclusive research perspective

### Justification 2: sex was not considered because …..

"All animals analysed were P3 or younger, thus no sex determination was attempted. Analyses are thought to include animals of both sexes at approximately equal proportions."

Assessment options:

- **Green** – design was appropriate from a sex inclusive research perspective
- **Amber** – research study raises some caution statements due to risk in the design/analysis
- **Red** – design was not appropriate from a sex inclusive research perspective

### Justification 3: single sex design because …..

"Because collecting vaginal smears to control oestrous cycle in females would add an extra layer of stress to our experiments, we performed the experiments in male pups only. "

Assessment options:

- **Green** – design was appropriate from a sex inclusive research perspective
- **Amber** – research study raises some caution statements due to risk in the design/analysis
- **Red** – design was not appropriate from a sex inclusive research perspective

### Justification 4: single sex design because …..

"We chose to use male mice, as female mice tend to have twice the levels of circulating CORT as males, and these levels may shift in response to stage of the oestrus cycle "

Assessment options:

- **Green** – design was appropriate from a sex inclusive research perspective
- **Amber** – research study raises some caution statements due to risk in the design/analysis
- **Red** – design was not appropriate from a sex inclusive research perspective

### Justification 5: Some studies had a single sex design because …..

"[we only used male mice as Foxp3Sf is X-linked]. The transfer experiments were with male mice as the Foxp3Sf Th9 cells were obtained from male mice and could not be transferred to females due to risk of rejection"

Assessment options:

- **Green** – design was appropriate from a sex inclusive research perspective
- **Amber** – research study raises some caution statements due to risk in the design/analysis
- **Red** – design was not appropriate from a sex inclusive research perspective

7

### Conclusions

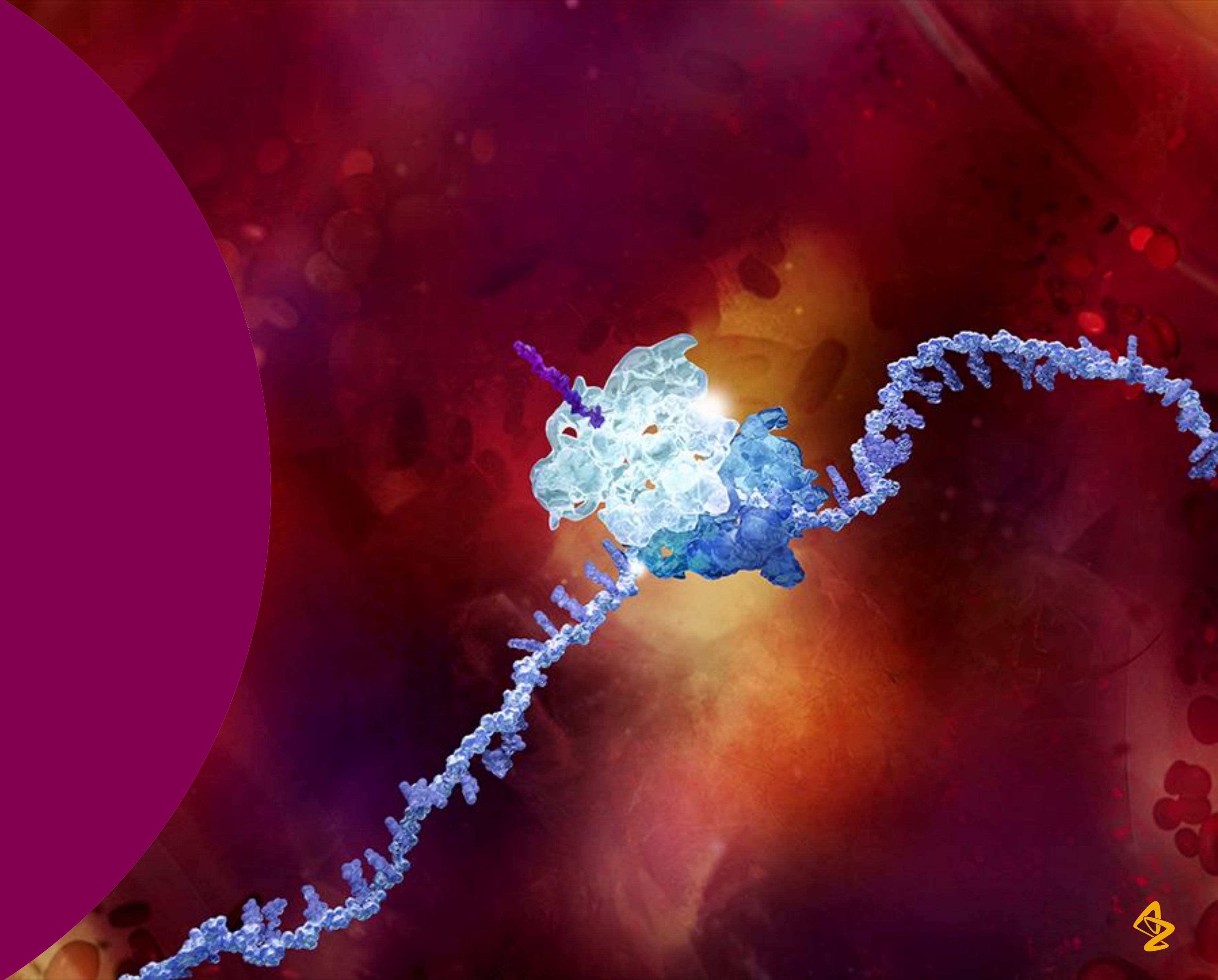

### Conclusions

- Research suggests that sex is a significant source of variation for both *in vivo* and *in vitro*.
- Improving translation requires us to embrace variation.  Sex is binary and is an easy first step to improve generalisability.
- However, sex bias is culturally embedded in our research pipelines, impacting the reporting, design, and  analysis.
- Many of the barriers are misconceptions.
- Including both sexes, is not at odds with a reduction mindset.
- The expectations is changing.  We need to work out how to consider sex mindfully and effectively.

### Acknowledgements

Neil Reavey
Natasha Karp
Ben Phillips
Timo Haschler

**Decision Tree development:**

Manuel Berdoy – University of Oxford
Lillian Hunt – Wellcome Trust
Maggy Jennings – RSPCA
Angela Kerton the Learning Curve Ltd
Matt Leach – Newcastle University
Jordi Lopez-Tremoleda - Queen Mary Uni of London
Jon Gledhill – Newcastle University
Esther J. Pearl – NC3Rs
Nathalie Percie du Sert – NC3Rs
Penny Reynolds – University of Florida
Kathy Ryder – Department of Health, Northern Ireland
Clare Stanford – UCL
Sara Wells – MRC Harwell
Lucy Whitfield – OWL Vets Ltd

**Prevalence of sexual dimorphism**

IMPC

UCL: Richard Mott, Judith Mank

Tel Aviv University: Ruth Heller, Shay Yaccoby, Yoav Benjamini

EMBL-EBI: Jeremy Mason



**Confidentiality Notice**

This file is private and may contain confidential and proprietary information. If you have received this file in error, please notify us and remove it from your system and note that you must not copy, distribute or take any action in reliance on it. Any unauthorized use or disclosure of the contents of this file is not permitted and may be unlawful. AstraZeneca PLC, 1 Francis Crick Avenue, Cambridge Biomedical Campus, Cambridge, CB2 0AA, UK, T: +44(0)203 749 5000, www.astrazeneca.com
