## Supplementary material 1 - WCP survey for "Addressing cultural and knowledge barriers to enable preclinical sex inclusive research": Supplementary material 1 - WCP survey.pdf

### Exploring researchers' perspective on sex inclusive in vivo research

If you have any questions about this survey, please find a survey facilitator or contact Prof. Amrita Ahluwalia or Dr. Jonathan Ho

* Required

1. By leaving your initials below, you acknowledge that you have had the opportunity to read the survey information sheet and consent form and have the research study explained. You have had the opportunity to ask questions about the research study, and your questions have been answered. You understand that if you are uncomfortable with any question, you may skip it. *

2. Does your current research use animal models to investigate your disease of interest? *

- ◯ Yes
- ◯ No

3. Does your research focus on a disease, or phenomena, that is only, or predominantly, seen in one sex? *

- ◯ Yes
- ◯ Sometimes
- ◯ No

4. How often are you involved or can influence the planning of experiments involving animals *

- ◯ Never
- ◯ Rarely
- ◯ Sometimes
- ◯ Often
- ◯ Always

5. In the past, how much statistical training have you received?

- ◯ No training
- ◯ Primarily informal/practical training (e.g., books, videos, discussion)
- ◯ 1-2 formal courses (e.g., Coursera, University class)
- ◯ More than 2 courses or a statistical degree

6. How familiar are you with factorial experimental designs (please consider your general knowledge and application of the technique)?

- ◯ Not at all familiar
- ◯ Not very familiar
- ◯ Somewhat familiar
- ◯ Familiar
- ◯ Extremely familiar

7. Thinking over the last 5 years, how often have you incorporated both sexes into your experiment while studying an intervention (for example a drug treatment)?

- ◯ Never: 0% of the time
- ◯ Rarely: 1-25% of the time
- ◯ Sometimes: 26-50% of the time
- ◯ Often: 51-75% of the time
- ◯ Almost always: 76-100% of the time

8. Select which of the following prevents you from including both sexes in future experiments in addition to the intervention of interest (choose **all** that apply):

- [ ] Not relevant to the research question
- [ ] Welfare issues
- [ ] Sample size concerns
- [ ] Female animals are more variable
- [ ] Complexity of experimental design
- [ ] Availability of sample/test material
- [ ] Data analysis concerns
- [ ] Cost
- [ ] Model behavior may be different in the other sex
- [ ] Male animals are more likely to fight and may lead to premature euthanasia
- [ ] Other

**Over the next few questions, answer based on your current thinking about how incorporating sex into an in vivo experimental design may affect the overall design, data or statistical analysis resulting from that study.**

9. Do you think inclusion of both sexes requires doubling a study's sample size?

- ◯ Yes
- ◯ Sometimes
- ◯ No
- ◯ Don't know

10. Do you think sex influences data variability, therefore when you include both sexes, more animals are needed?

- ◯ Yes
- ◯ Sometimes
- ◯ No
- ◯ Don't know

11. When analyzing *in vivo* data collected from both sexes, do you think sex should be included in the statistical model?

- ◯ Yes
- ◯ Sometimes
- ◯ No
- ◯ Don't know

12. When analyzing *in vivo* data, do you think data from the two sexes should be pooled (combined) for an intervention into a single group for the analysis?

- ◯ Yes
- ◯ Sometimes
- ◯ No
- ◯ Don't know

13. When analyzing *in vivo* data collected from both sexes, do you think the analysis should be run independently for each sex through separate statistical tests?

- ◯ Yes
- ◯ Sometimes
- ◯ No
- ◯ Don't know

27. Select which of the following prevents you from **including both sexes** in future experiments in addition to the intervention of interest (choose **all** that apply):

- [ ] Not relevant to the research question
- [ ] Welfare issues
- [ ] Sample size concerns
- [ ] Female animals are more variable
- [ ] Complexity of experimental design
- [ ] Availability of sample/test material
- [ ] Data analysis concerns
- [ ] Cost
- [ ] Experiment would take longer
- [ ] Other

28. Select which of the following you believe are the advantages, if any, to **including both sexes in an in vivo experimental design**? (choose **all** that apply):

- [ ] Translatability
- [ ] Reproducibility
- [ ] Understanding sex differences
- [ ] Efficient use of all animals from breeding
- [ ] Animal welfare
- [ ] 3Rs – Reduction
- [ ] Other

29. Do you have any other thoughts or comments you would like to provide regarding **including both sexes in an in vivo experimental design**?

30. Are you completing this survey as part of attending

- ◯ The WCP general conference
- ◯ Symposium - The importance of interrogating sex differences in cardiovascular physiology and disease (July 4)
- ◯ Workshop - Best practice for sex inclusive research (July 5)

31. Which of the following have you already attended

Please select at most 2 options.

- ☐ Symposium - The importance of interrogating sex differences in cardiovascular disease (July 4th)
- ☐ Workshop - Best practice for sex inclusive research (July 5th)
- ☐ None of the above

32. What is your age?

Number must be between 18 ~ 110

33. Which of the following best describes your gender? (If you would prefer to self-describe, please do so in the free-text box.)

- ◯ Man
- ◯ Non-binary
- ◯ Woman
- ◯ Prefer not to say
- ◯ Other

34. What geographic region is your primary job located in?

- Africa
- North America
- Latin America
- Asia
- Australia
- Europe
- Middle East

35. How many years have you worked with animals in research?

Number must be between 0 ~ 70

36. What type of institution do you currently work for in your primary role?

- Academic institution
- Contract research organization
- Pharmaceutical company
- Biotechnology company
- Non-profit
- Publisher (e.g., scientific journal or book)
- Scientific measurement company (e.g., develop assays or biological sample analysis kits)
- Consultant
- Other

37. What is the primary type of *in vivo* research in which you are involved (if several, chose the option you identify most with)?

- Basic biological investigation
- Discovery (e.g., target identification)
- Drug development/Lead optimization
- Production quality and manufacturing
- Research support (e.g., statisticians, ethical review board, or animal technician)
- Safety pharmacology/preclinical safety/toxicology
- Other

38. What is your highest level of education?

- Doctoral degree (PhD, veterinary degree, other)
- Master's degree
- Bachelor's degree (4 years at college/university)
- Associate or vocational qualification (2 years at college/university)
- High school diploma, GED, A level, or equivalent
- Other

39. If you'd like to be entered into the draw for a £50 Amazon gift card, please leave your email address below

This content is neither created nor endorsed by Microsoft. The data you submit will be sent to the form owner.

Microsoft Forms
